# Systematic mapping of chemoreceptor specificities for *Pseudomonas aeruginosa*

**DOI:** 10.1101/2023.04.04.535651

**Authors:** Wenhao Xu, Jean Paul Cerna-Vargas, Ana Tajuelo, Andrea Lozano Montoya, Melissa Kivoloka, Nicolas Krink, Elizabet Monteagudo-Cascales, Miguel A. Matilla, Tino Krell, Victor Sourjik

## Abstract

The chemotaxis network, one of the most prominent prokaryotic sensory systems, is present in most motile bacteria and archaea. Although the conserved signaling core of the network is well characterized, ligand specificities of a large majority of diverse chemoreceptors encoded in bacterial genomes remain unknown. Here we performed a systematic identification and characterization of new chemoeffectors for the opportunistic pathogen *Pseudomonas aeruginosa*, which has 26 chemoreceptors possessing most of the common types of ligand binding domains. By performing capillary chemotaxis assays for a library of growth-promoting compounds, we first identified a number of novel chemoattractants of varying strength. We subsequently mapped specificities of these ligands by performing Förster resonance energy transfer (FRET) and microfluidic measurements for hybrids containing ligand binding domains of *P. aeruginosa* chemoreceptors and the signaling domain of the *Escherichia coli* Tar receptor. Direct binding of ligands to chemoreceptors was further confirmed *in vitro* using thermal shift assay and microcalorimetry. Altogether, the combination of methods enabled us to assign several new attractants, including methyl 4-aminobutyrate, 5-aminovalerate, L-ornithine, 2-phenylethylamine and tyramine, to previously characterized chemoreceptors and to annotate a novel purine-specific receptor PctP. Our screening strategy could be applied for the systematic characterization of unknown sensory domains in a wide range of bacterial species.

**Importance:** Chemotaxis of motile bacteria has multiple physiological functions. It enables bacteria to locate optimal ecological niches, mediates collective behaviors, and can play an important role in infection. These multiple functions largely depend on ligand specificities of chemoreceptors, and the number and identities of chemoreceptors show high diversity between organisms. Similar diversity is observed for the spectra of chemoeffectors, which include not only chemicals of high metabolic value but also bacterial, plant and animal signaling molecules. However, the systematic identification of chemoeffectors and their mapping to specific chemoreceptors remains a challenge. Here, we combined several *in vivo* and *in vitro* approaches to establish a systematic screening strategy for the identification of receptor ligands, and we applied it to identify a number of new physiologically relevant chemoeffectors for the important opportunistic human pathogen *P. aeruginosa*. This strategy can be equally applicable to map specificities of sensory domains from a wide variety of receptor types and bacteria.

## Introduction

Most bacteria have evolved the ability to detect a wide range of environmental signals to survive and grow under rapidly changing conditions. One of the most prominent prokaryotic sensory systems is the chemotaxis network that controls motility (1, 2). Chemotaxis has multiple important functions in bacterial physiology, dependent on the lifestyle and ecological niche, enabling bacteria to move towards optimal growth environments but also mediating collective behaviors and interactions with eukaryotic hosts (3, 4).

Such variety of chemotaxis-mediated functions is primarily ensured by the diversity of bacterial chemoreceptors, also called methyl-accepting chemotaxis proteins (MCPs) (5). While the core of the signaling pathway is conserved among bacteria, the number and specificity of chemoreceptors is highly variable and strain-specific (6). The reported repertoire of signals recognized by chemoreceptors across bacterial species includes proteinogenic amino acids (7, 8), polyamines (9), quaternary amines (10), nucleobases and their derivatives (11, 12), organic acids (13, 14), sugars (15), but also inorganic ions (16, 17), pH (18–20) and temperature (21, 22). Nevertheless, the signal specificity remains unknown for the absolute majority of chemoreceptors.

The paradigmatic model system of *Escherichia coli* chemotaxis consists of a single pathway, controlled by four transmembrane chemoreceptors and one aerotaxis receptor, and it includes six cytoplasmic signaling proteins: a histidine kinase CheA, an adaptor CheW, a response regulator CheY, a methyltransferase CheR, a methylesterase CheB, and a phosphatase CheZ (2). Typically, chemotactic stimuli modulate the autophosphorylation activity of CheA, which is inhibited by attractants and stimulated by repellents, subsequently altering the transphosphorylation of CheY. The phosphorylated CheY binds to the flagellar motor resulting in a change in the direction of flagellar rotation, ultimately causing a chemotactic response. CheZ is responsible for the dephosphorylation of CheY. After the initial pathway response, an adaptation system composed of CheR and CheB adjusts the level of receptor methylation on several specific glutamyl residues, providing negative feedback to the kinase activity, which ensures adaptation of cells to persisting stimulation. Although most of the chemotaxis proteins found in *E. coli* are conserved across bacterial chemotaxis pathways, most bacteria have more complex chemosensory pathways, possessing additional chemotaxis proteins and chemoreceptors, alternative adaptation and signal termination strategies (23–25).

Canonical chemoreceptors can be separated in three functional domains: a periplasmic ligand binding domain (LBD), a signal conversion HAMP domain, and a cytoplasmic signaling domain that interacts with the autokinase CheA (5). Analyses of sequenced bacterial genomes revealed that bacterial chemoreceptors employ more than 80 different types of LBDs (6). In contrast, all *E. coli* transmembrane chemoreceptors possess the same four-helix bundle (4HB) type of LBD. Thus, although *E. coli* chemotaxis signaling pathway is one of the simplest and best understood, it does not represent the diversity of bacterial sensory capabilities.

*Pseudomonas aeruginosa* is among the most important human pathogens, causing the death of more than half a million people annually (26), and it is also one of the most well-studied alternative models for chemotaxis (27, 28). The chemoreceptor repertoire of the *P. aeruginosa* model strain PAO1 has 26 chemoreceptors containing 12 different LBD types that feed into four different chemosensory pathways, of which 23 chemoreceptors were predicted to stimulate the genuine F6-type chemotaxis pathway that controls swimming motility. Other three receptors McpB/Aer2, WspA, and PilJ are involved in an F7-type pathway of unknown function, an alternative cellular function pathway that mediates c-di-GMP synthesis, and type IV pili chemosensory pathway that is associated with twitching motility, respectively. The latter two pathways were found to perform mechano- and surface sensing rather than chemosensing (29–31). Of the 18 chemoreceptors with a periplasmic LBD that belong to the F6 pathway (28), ten have been functionality annotated (Table 1). Ligands of the other eight chemoreceptors involved in chemotactic behaviors are yet to be characterized, and given the variety of ligands that are typically sensed by a single LBD, even the annotated LBDs are likely to possess additional ligand specificities.

**Table 1.**
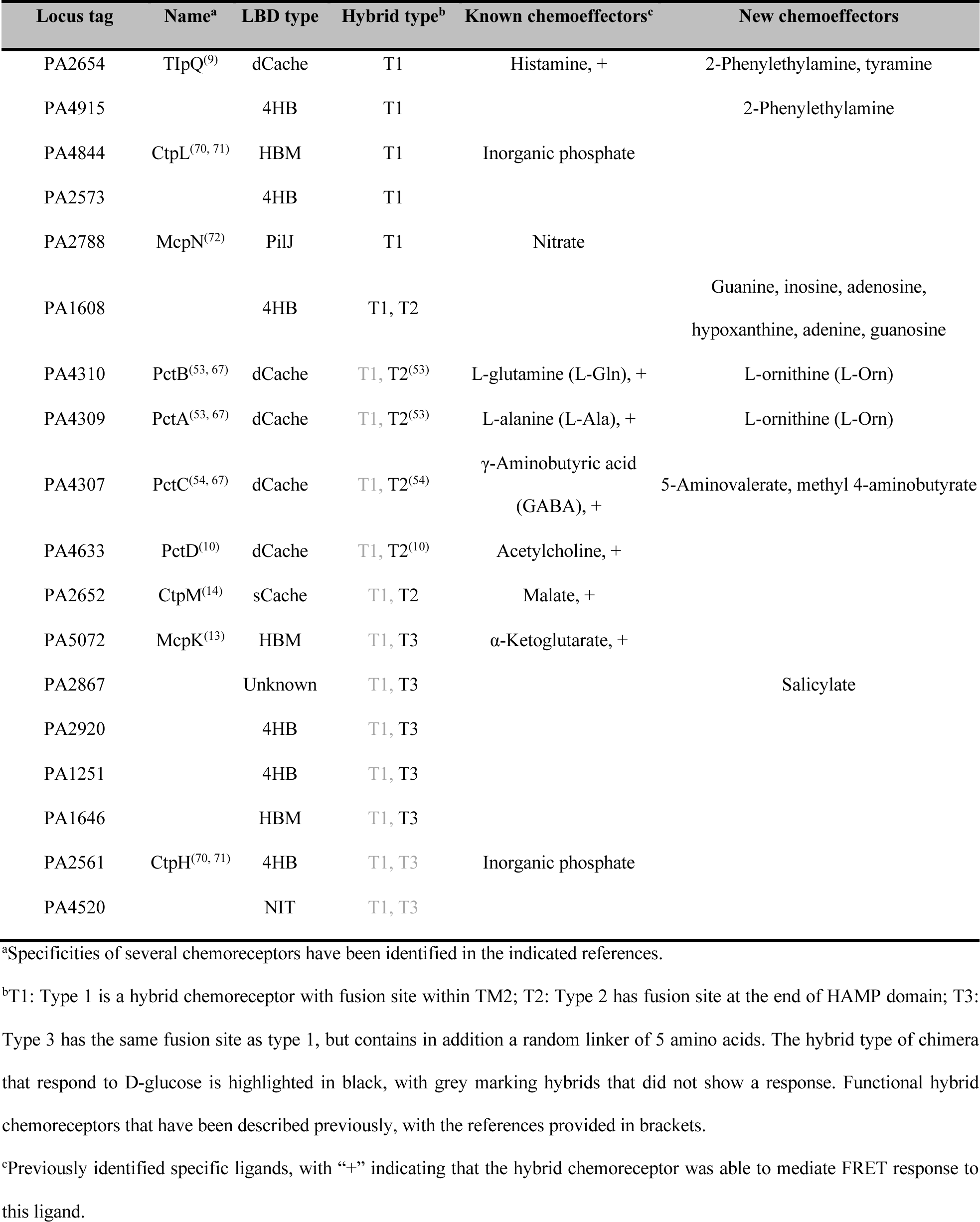
Summary of all the chimeras constructed and new chemoeffectors identified in this work.

| Locus tag | Name<sup>a</sup> | LBD type | Hybrid type<sup>b</sup> | Known chemoeffectors<sup>c</sup> | New chemoeffectors |
| --- | --- | --- | --- | --- | --- |
| PA2654 | TlpQ<sup>(9)</sup> | dCache | T1 | Histamine, + | 2-Phenylethylamine, tyramine |
| PA4915 |  | 4HB | T1 |  | 2-Phenylethylamine |
| PA4844 | CtpL<sup>(70, 71)</sup> | HBM | T1 | Inorganic phosphate |  |
| PA2573 |  | 4HB | T1 |  |  |
| PA2788 | McpN<sup>(72)</sup> | PilJ | T1 | Nitrate |  |
| PA1608 |  | 4HB | T1, T2 |  | Guanine, inosine, adenosine, hypoxanthine, adenine, guanosine |
| PA4310 | PctB<sup>(53, 67)</sup> | dCache | T1, T2<sup>(53)</sup> | L-glutamine (L-Gln), + | L-ornithine (L-Orn) |
| PA4309 | PctA<sup>(53, 67)</sup> | dCache | T1, T2<sup>(53)</sup> | L-alanine (L-Ala), + | L-ornithine (L-Orn) |
| PA4307 | PctC<sup>(54, 67)</sup> | dCache | T1, T2<sup>(54)</sup> | γ-Aminobutyric acid (GABA), + | 5-Aminovalerate, methyl 4-aminobutyrate |
| PA4633 | PctD<sup>(10)</sup> | dCache | T1, T2<sup>(10)</sup> | Acetylcholine, + |  |
| PA2652 | CtpM<sup>(14)</sup> | sCache | T1, T2 | Malate, + |  |
| PA5072 | McpK<sup>(13)</sup> | HBM | T1, T3 | α-Ketoglutarate, + | |
| PA2867 |  | Unknown | T1, T3 |  | Salicylate |
| PA2920 |  | 4HB | T1, T3 |  |  |
| PA1251 |  | 4HB | T1, T3 |  |  |
| PA1646 |  | HBM | T1, T3 |  |  |
| PA2561 | CtpH<sup>(70, 71)</sup> | 4HB | T1, T3 | Inorganic phosphate |  |
| PA4520 |  | NIT | T1, T3 |  |  |

<sup>a</sup>Specificities of several chemoreceptors have been identified in the indicated references.
<sup>b</sup>T1: Type 1 is a hybrid chemoreceptor with fusion site within TM2; T2: Type 2 has fusion site at the end of HAMP domain; T3: Type 3 has the same fusion site as type 1, but contains in addition a random linker of 5 amino acids. The hybrid type of chimera that respond to D-glucose is highlighted in black, with grey marking hybrids that did not show a response. Functional hybrid chemoreceptors that have been described previously, with the references provided in brackets.
<sup>c</sup>Previously identified specific ligands, with “+” indicating that the hybrid chemoreceptor was able to mediate FRET response to this ligand.

Several experimental approaches have been developed to systematically characterize the specificities of uncharacterized chemoreceptors. The quantitative capillary chemotaxis assay is a traditional method for identifying bacterial chemoeffectors. However, because of differences in motility and physiology, the experimental conditions for the capillary assay need to be established for each individual bacterial strain, which complicates its general application. Moreover, assignment of identified ligands to specific chemoreceptors typically requires the construction of strains with deletions of individual receptor genes, and it is complicated by the frequent functional redundancy of multiple chemoreceptors. Alternatively, biochemical assays can be used for ligand identification *in vitro* (32). The thermal shift assay (TSA; alternatively called differential scanning fluorimetry, DSF) can be applied to characterize binding of chemical compounds to LBDs in high-throughput screens, but it is prone to yield false-positive results. Isothermal titration calorimetry (ITC) is, in contrast, an accurate but low-throughput method to measure ligand binding (33). Although a combination of these *in vitro* methods has proven to be very powerful for ligand identification (34), their application is limited to the LBD that can be purified and to compounds with high-affinity binding.

A complementary strategy for identification of receptor ligands relies on the construction of chimeric receptors that combine an LBD of interest with the well-characterized output domain, such as that of the *E. coli* Tar receptor. Both, chemoreceptor-chemoreceptor hybrids (35) and chemoreceptor-histidine kinase hybrids (36) that enable an *in vivo* readout of signaling response have been recently used to annotate unknown sensory functions. Here, we constructed a library of most *P. aeruginosa* LBDs fused to the signaling domain of the *E. coli* chemoreceptor Tar, and we used this library in combination with *in vivo* and *in vitro* assays to identify several novel physiologically relevant chemoeffectors and to assign them to specific *P. aeruginosa* chemoreceptors. Overall, our screening strategy allowed us to expand the list of known chemoreceptor specificities for *P. aeruginosa*, and a similar approach should be applicable to chemoreceptors from other bacteria and even to other types of receptors with a periplasmic LBD.

## Results

### High-throughput screening for putative chemoeffectors in *P. aeruginosa*

To identify potential chemoeffectors for *P. aeruginosa*, we first screened chemical compounds from a large library of metabolites. Since several studies have shown correlation between a metabolic value of a compound and its potency as a chemoeffector (8, 37), we first used a growth assay to test effects of chemical compounds from three plates of the commercial Biolog compound arrays (PM1, PM2A and PM3B). This growth assay indeed showed that 202 compounds could be utilized as either carbon or nitrogen sources to support *P. aeruginosa* growth (Table S1). Out of those, we selected 30 compounds from Biolog compound arrays and additional 9 compounds that have been characterized as the specific ligands in other strains of *Pseudomonas* spp. These 39 compounds have not yet been characterized as chemoattractants for *P. aeruginosa*, indicating that at least some of them are likely to be novel ligands for *P. aeruginosa* chemoreceptors.

We next investigated the chemotactic responses of *P. aeruginosa* to these potential ligands by performing quantitative capillary chemotaxis assays (Fig. 1). Five compounds were able to highly efficiently attract bacteria, with the strongest response being observed for methyl 4-aminobutyrate, followed by 5-aminovalerate, ethanolamine (EA), L-ornithine, and 2-phenylethylamine (PEA). Other seven compounds, guanine, glutarate, tricarballylate, niacinate, succinate, fumarate and formate, acted as chemoattractants of intermediate strength. Of note, low concentration of guanine (10 µM instead of 1 mM) was used in this assay due to its poor solubility. The remaining compounds mediated only weak or no chemotaxis, despite being nutrients.

**Fig. 1.**
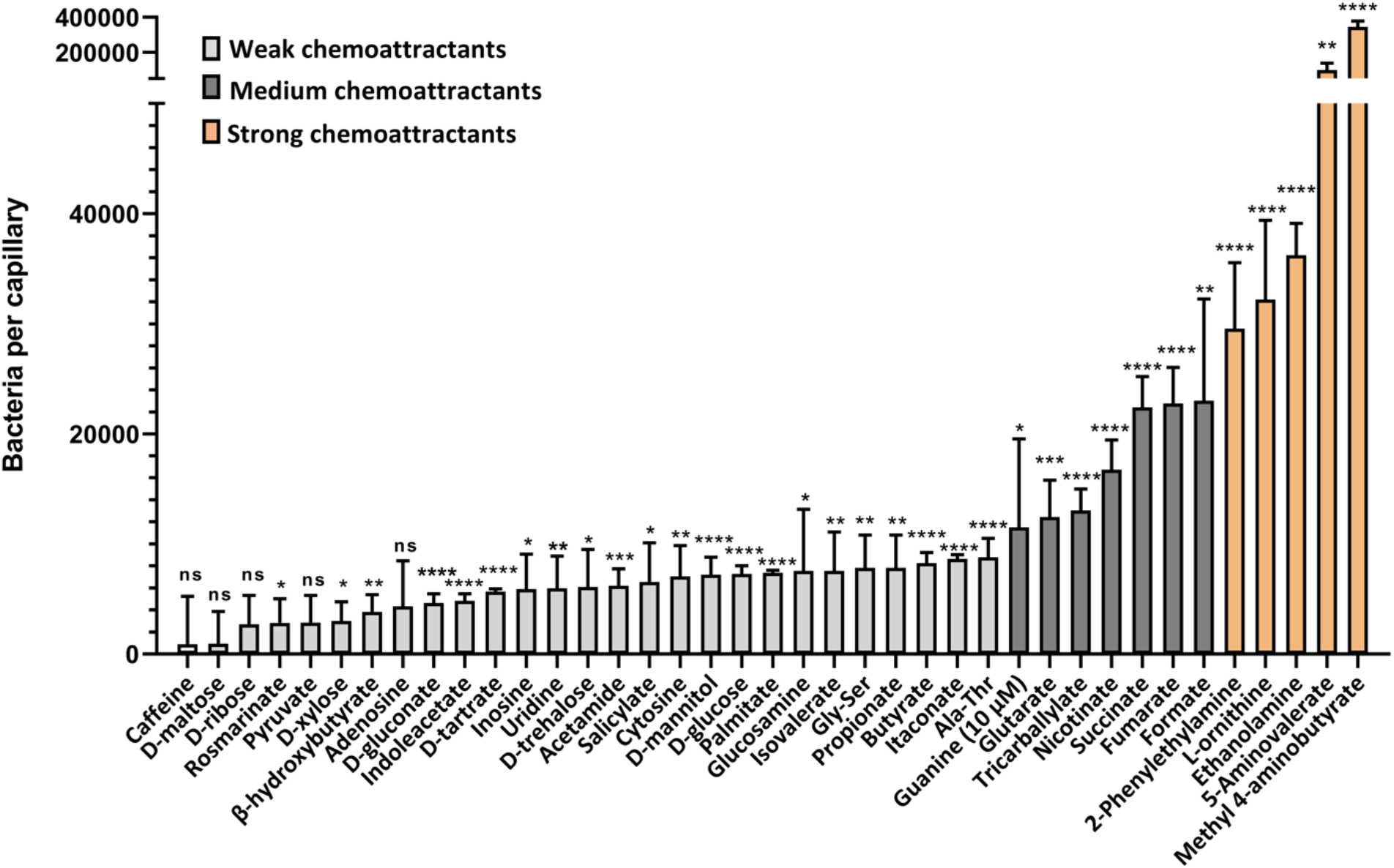
Chemotaxis of *P. aeruginosa* PAO1 toward potential chemoeffectors. Accumulation of bacteria (WT-Washington) in capillaries containing 1 mM of indicated chemical compound in the chemotaxis buffer, except guanine that was used at 10 µM due to poor solubility. Strong chemoattractants with >30,000 cells per capillary are shown in orange; medium chemoattractants with 10,000 - 30,000 cells per capillary are shown in dark grey and the remaining chemical compounds are shown in light grey. All data have been corrected by the number (7,825 ± 623) of bacteria in buffer-containing capillaries. The means and standard deviations of four biological replicates each conducted in triplicate are shown. Error bars indicate the mean ± standard deviations. Significance of difference was statistically significant differences from buffer-containing capillaries, assessed using an unpaired Student’s t-test, are indicated by asterisks (ns: non-significant, *p ≤ 0.05, **p ≤ 0.01, ***p ≤ 0.001, and ****p ≤ 0.0001).

### Construction of chimeras for 18 transmembrane chemoreceptors in *P. aeruginosa*

To investigate the specificities of these unassigned chemoeffectors, we focused on the 18 transmembrane chemoreceptors that belong to the F6 chemotaxis pathway of *P. aeruginosa* (Table 1). To increase the probability of obtaining functional hybrid chemoreceptor for each candidate, we applied three previously described construction strategies with different fusion sites: within the transmembrane (TM) helix 2, after the HAMP domain, and within the TM2 helix with a 5 amino acids random linker (17, 35). The resulting hybrids of type 1 and type 3 connect the extracellular sensory domain, the TM1 helix, and part of the TM2 helix from *P. aeruginosa* chemoreceptor to the rest of the TM2 helix, the HAMP domain and the cytoplasmic signaling domain of *E. coli* chemoreceptor Tar (Fig. 2A). The fusion site of type 2 chimera is located in the junction between the HAMP domain and cytoplasmic signaling domain, thus connecting the extracellular sensory domain, the two TM helices and the whole HAMP domain from *P. aeruginosa* chemoreceptor to the cytoplasmic signaling domain of Tar.

**Fig. 2.**
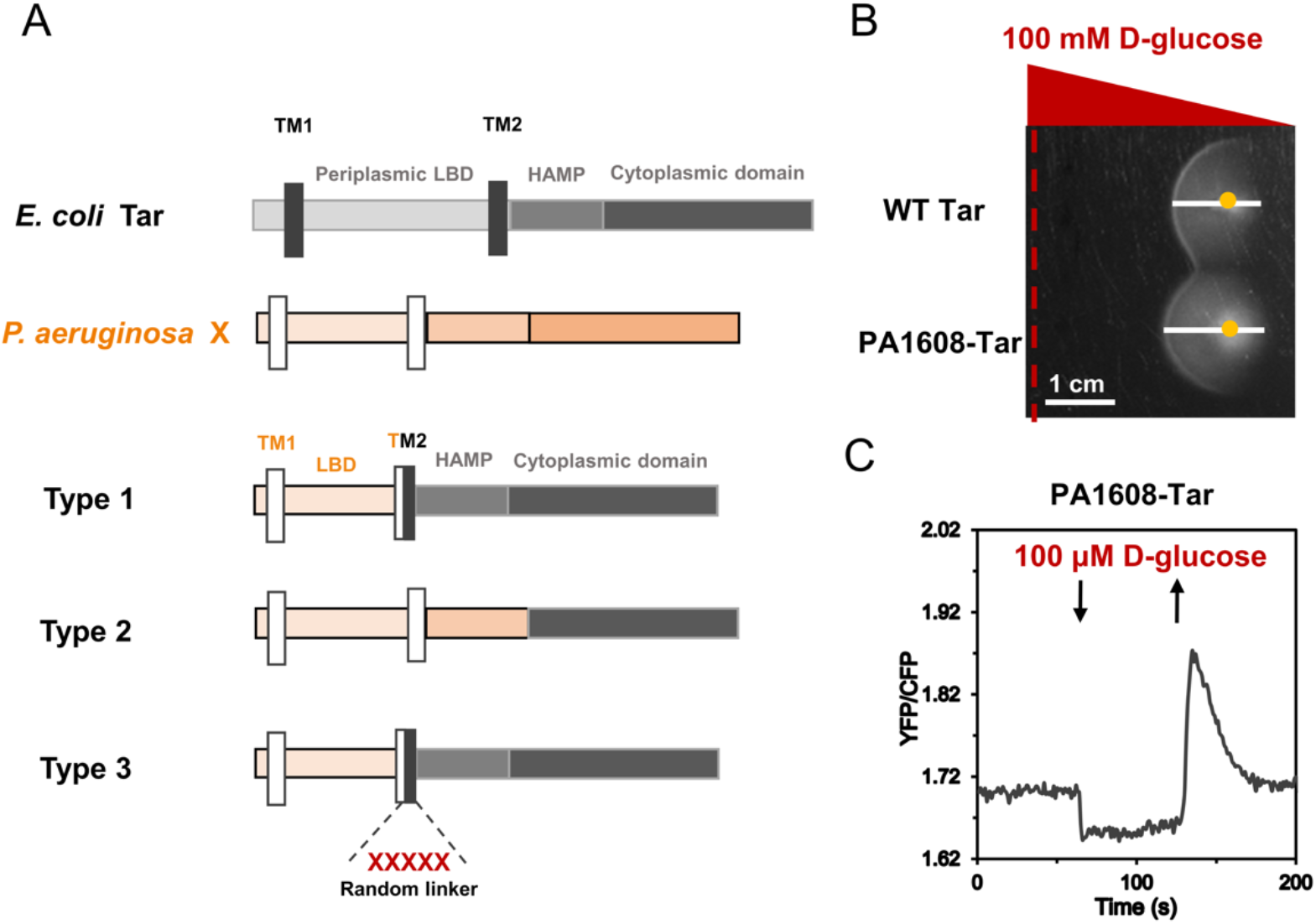
Construction and evaluation of receptor hybrids for transmembrane chemoreceptors of *P. aeruginosa* PAO1. (A) Schematic drawing of three types of hybrid chemoreceptors constructed in this work. Tar chemoreceptor from *E. coli* and target chemoreceptor (X) from *P. aeruginosa* each consist of three parts, periplasmic ligand binding domain, HAMP domain and cytoplasmic domain, shown in different colors. The two TM helices are shown by black and white rectangles in the Tar receptor and the target chemoreceptor (X), respectively. (B) D-glucose gradient plate assay used for the assessment of receptor functionality. The control was the receptorless *E. coli* cells expressing wildtype Tar as the sole receptor. Positions of the ligand source and inoculated cells are shown by a red dotted line and by a yellow dot, respectively. Spreading of cells toward or away from the ligand source is indicated by the white line (scale bar, 1 cm). (C) FRET measurement of the hybrid chemoreceptor response to D-glucose. Receptorless *E. coli* cells expressing the CheZ-CFP/CheY-YFP FRET pair and PA1608-Tar as the sole receptor responded to stepwise addition (down arrow) and subsequent removal (up arrow) of the 100 µM D-glucose. Similar to canonical chemoattractants, addition of D-glucose inhibits pathway activity and thus lowers the YFP/CFP fluorescence ratio.

We subsequently tested the functionality of these chimeras expressed in a receptorless *E. coli* strain. We first used soft-agar gradient plates, where chemotactic cells exhibit biased spreading in gradients of compounds that are established by diffusion (Fig. 2B). Subsequently, we performed Förster resonance energy transfer FRET measurements (Fig. 2C) that are based on the phosphorylation-dependent interaction between CheY fused to yellow fluorescent protein (CheY-YFP) and CheZ fused to cyan fluorescent protein (CheZ-CFP). This assay enables to monitor activity of the chemotaxis pathway by following changes in the ratio YFP/CFP fluorescence, which is proportional to CheA activity (38). Because ligand specificity for many tested LBDs is not known, D-glucose was routinely used as a non-specific chemoattractant to assess the activity of hybrids. Differently from conventional chemoattractants, D-glucose is sensed via the phosphotransferase system (PTS) which signals to the cytoplasmic part of the chemoreceptor (39–41). A response to D-glucose thus demonstrates that the receptor hybrid activates the pathway and is responsive to stimulation, but the functionality of extracellular sensory domain and signal transduction toward the cytoplasmic part of the receptor remains to be confirmed by a specific ligand. From the constructed hybrids, only PA2561-Tar and PA4520-Tar showed no response to D-glucose. The functionality of several hybrids with already characterized LBDs was further verified by measuring FRET responses to their specific ligands (Table 1). Collectively, 16 active hybrid chemoreceptors were constructed successfully, and 7 of them were confirmed to be functional.

### Screening of specific ligands for chemoreceptor chimeras using FRET

For these 16 functional hybrids, we conducted the one-by-one screening with potential ligands at a fixed concentration of 100 μM, using FRET measurements in *E. coli* strains expressing the indicated hybrid chemoreceptor as a sole chemoreceptor along with the CheZ-CFP/CheY-YFP FRET pair. A number of tested ligands elicited similar responses for all (or most) receptor hybrids, as well as for the full-length Tar (Table S2), indicating that these stimuli might be sensed by the cytoplasmic portion of Tar (2). Responses to other potential ligands were hybrid-specific, suggesting that their selectivity for a particular LBD (Table 1 and Table S2).

As an example, a hybrid that contains a 4HB-type LBD of PA1608 with unknown function showed not only responses to D-glucose but also similarly strong responses to inosine and guanine (Fig. 3A). These compounds did not stimulate Tar (Fig. S1A-C), suggesting that the extracellular sensor domain of PA1608 might be specific for inosine and guanine (see below). Another example is the hybrid with the LBD of PctB, a well-known amino acid chemoreceptor, where the response to L-glutamine was used to confirm its functionality. Different from Tar (Fig. S1B), PctB-Tar was also able to produce strong FRET response upon exposure to L-ornithine (Fig. 3B). Other identified ligands are summarized in Table 1 and Table S2. Besides purines and L-ornithine, these included methyl 4-aminobutyrate and 5-aminovalerate (specific to PctC) and salicylate (specific to PA2867).

**Fig. 3.**
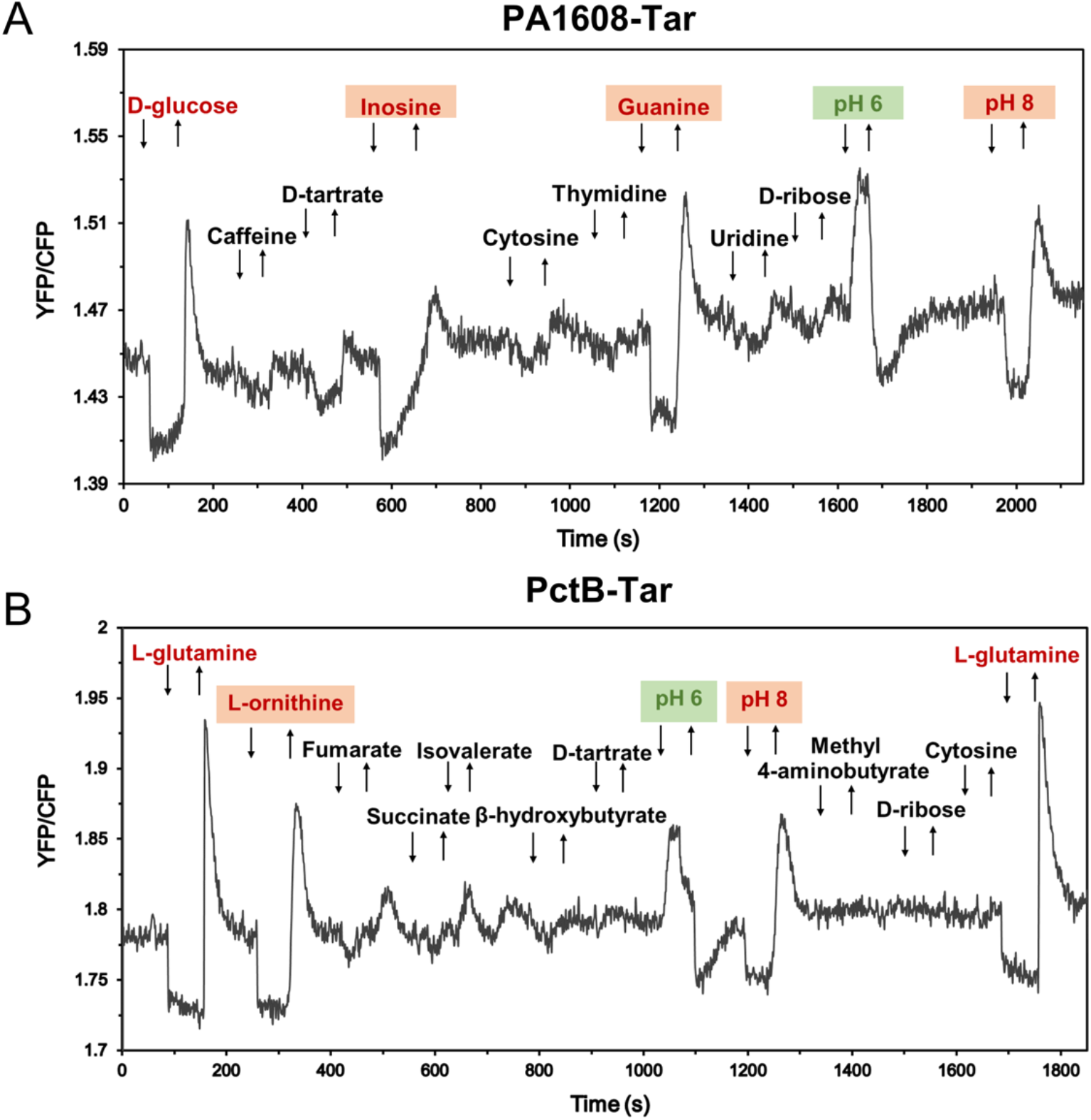
High-throughput screening of unassigned potential ligands for chemoreceptor chimeras with FRET. Examples of FRET measurements of receptorless *E. coli* FRET strain expressing PA1608-Tar (A) or PctB-Tar (B) as the sole receptor, responding to the stepwise addition (down arrow) and subsequent removal (up arrow) of indicated chemical compounds at concentration of 100 µM. Because of lack of previously characterized ligand for PA1608, D-glucose was used as a positive control for PA1608-Tar hybrid during FRET measurements to confirm receptor functionality. L-glutamine was used as a positive ligand for PctB-Tar hybrid. The chemoattractants are indicated in red and the chemorepellents were shown in green; newly identified chemoeffectors are shaded.

In addition to molecular compounds, we tested the response to external pH that is also an important stimulus for bacterial chemotaxis receptors. Furthermore, several pH-active compounds present in the Biolog compound arrays used in our initial screening altered the pH of the analysis buffer. We observed that several hybrids including PctA-Tar, PctB-Tar, PA1608-Tar and TlpQ-Tar mediated a repellent response to acidic pH 6.0 (Fig. 3), whereas PctC-Tar and PA1646-Tar showed an attractant response to acidic pH (Table S2). This indicates that, similar to that of *E. coli* (20), *P. aeruginosa* might exhibit bidirectional navigation in pH gradients in order to accumulate toward optimal pH.

### Characterization of novel chemoeffectors for amino acid chemoreceptors PctA, PctB and PctC

In general, whenever clear responses to a particular ligand were observed in the micromolar concentration range in the *E. coli* FRET strain expressing chimera as a sole receptor, whereas the strain expressing wildtype Tar showed no response, it was seen as evidence that ligand is recognized by the respective sensory domain. The FRET results showed that, besides their known amino acid ligands, PctA-Tar and PctB-Tar were responsive to L-ornithine (Fig. 4A-C), whereas no response could be observed for Tar (Fig. S2A). This observation is consistent with a recent report, although direct binding of L-ornithine to these receptors was not demonstrated (42). We therefore performed ITC experiments using purified the LBDs of PctA and PctB, which confirmed binding of L-ornithine to both LBDs, with higher affinity (*K_D_* = 7.1 μM) for PctA than for PctB (*K_D_* = 559 μM) (Fig. 4D and E). This difference in affinity was consistent with the relative potency of these ligands as chemoeffectors for hybrids in *E. coli* (Fig. 4C), when taking into account the *in vivo* signal amplification by the *E. coli* chemotaxis system.

**Fig. 4.**
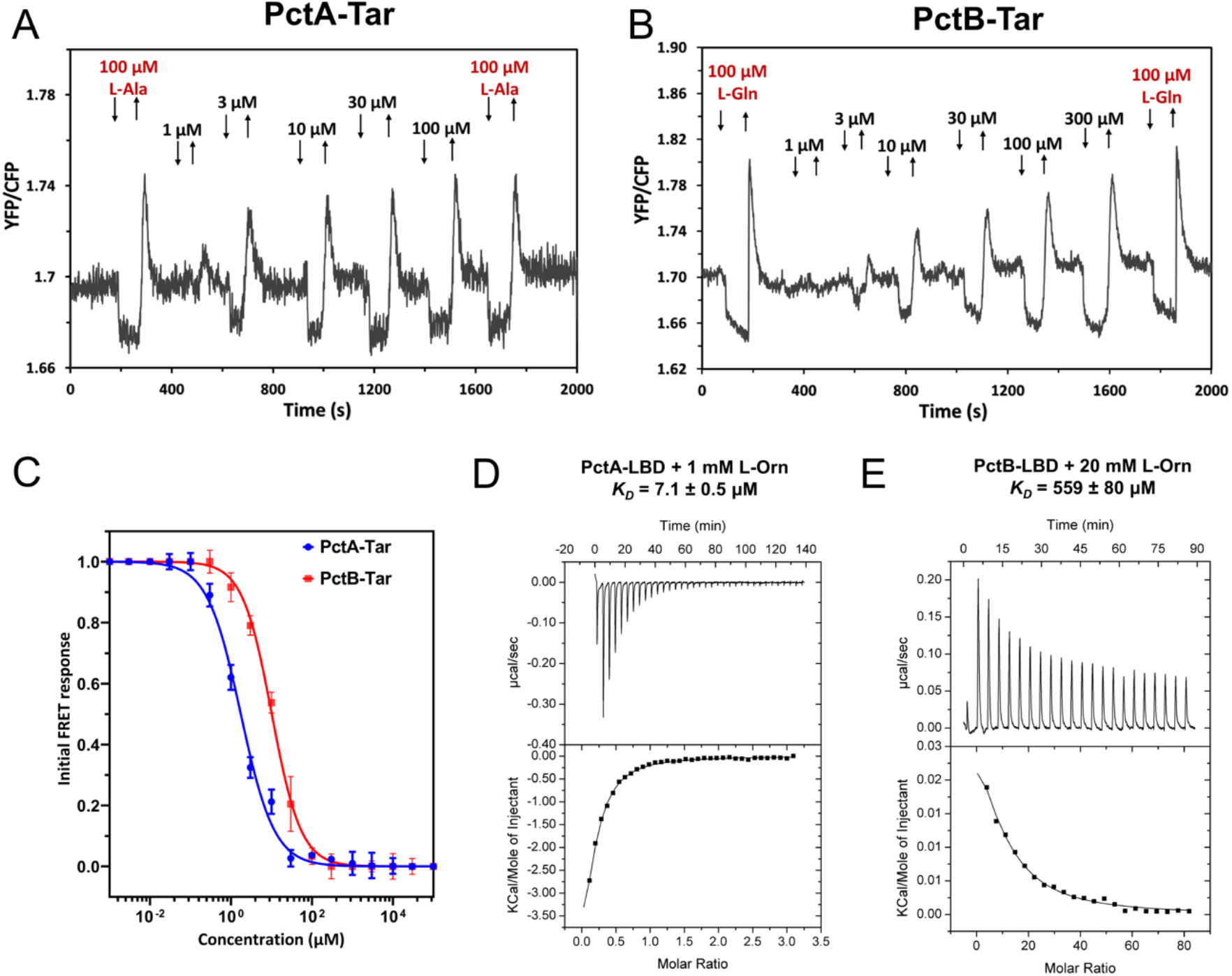
Characterization of L-ornithine sensing by PctA and PctB *in vivo* and *in vitro*. (A-C) FRET measurements of response to the indicated concentrations of L-ornithine (L-Orn) in *E. coli* expressing PctA-Tar (A) or PctB-Tar (B) as a sole receptor. The known ligands L-Ala (L-alanine) and L-Gln (L-glutamine) were used as positive controls for PctA-Tar and PctB-Tar, respectively. For corresponding dose-response curves (C), the amplitudes of the initial FRET response were calculated from changes in the ratio of YFP/CFP fluorescence after stimulation with indicated ligand concentrations and normalized to the saturated response. Error bars indicate the standard errors of three independent experiments; wherever invisible, error bars are smaller than the symbol size. Data were fitted using Hill equation, with the EC50 fit values being 1.8 ± 0.2 μM for PctA-Tar and 10.2 ± 0.8 μM for PctB-Tar. (D, E) Microcalorimetric titrations of PctA-LBD (D) and PctB-LBD (E) with L-Orn. The upper panels show raw titration data, and lower panels show integrated corrected peak areas of the titration data fit using the single-site binding model. Further experimental detail is provided in Table S4.

Similarly, the specificity of PctC-Tar response to methyl 4-aminobutyrate and 5-aminovalerate was confirmed by the dose-response FRET measurements (Fig. 5A-C and Fig. S2B-C). We further demonstrated direct binding of methyl 4-aminobutyrate (Fig. 5D) and 5-aminovalerate (Fig. 5E) to PctC-LBD by ITC, with dissociation constants of 129 and 28 μM, respectively. Again, these values were consistent with the higher sensitivity of PctC-Tar (lower EC50) to 5-aminovalerate in FRET measurements (Fig. 5C), taking in consideration the higher sensitivity of the *in vivo* response as discussed above. Finally, the relevance of the PctC-mediated response for chemotaxis of *P. aeruginosa* towards methyl 4-aminobutyrate and 5-aminovalerate was confirmed by the capillary chemotaxis assays, where the *pctC* deletion strain showed strongly reduced chemotaxis compared to the wildtype strain (Fig. 5F).

**Fig. 5.**
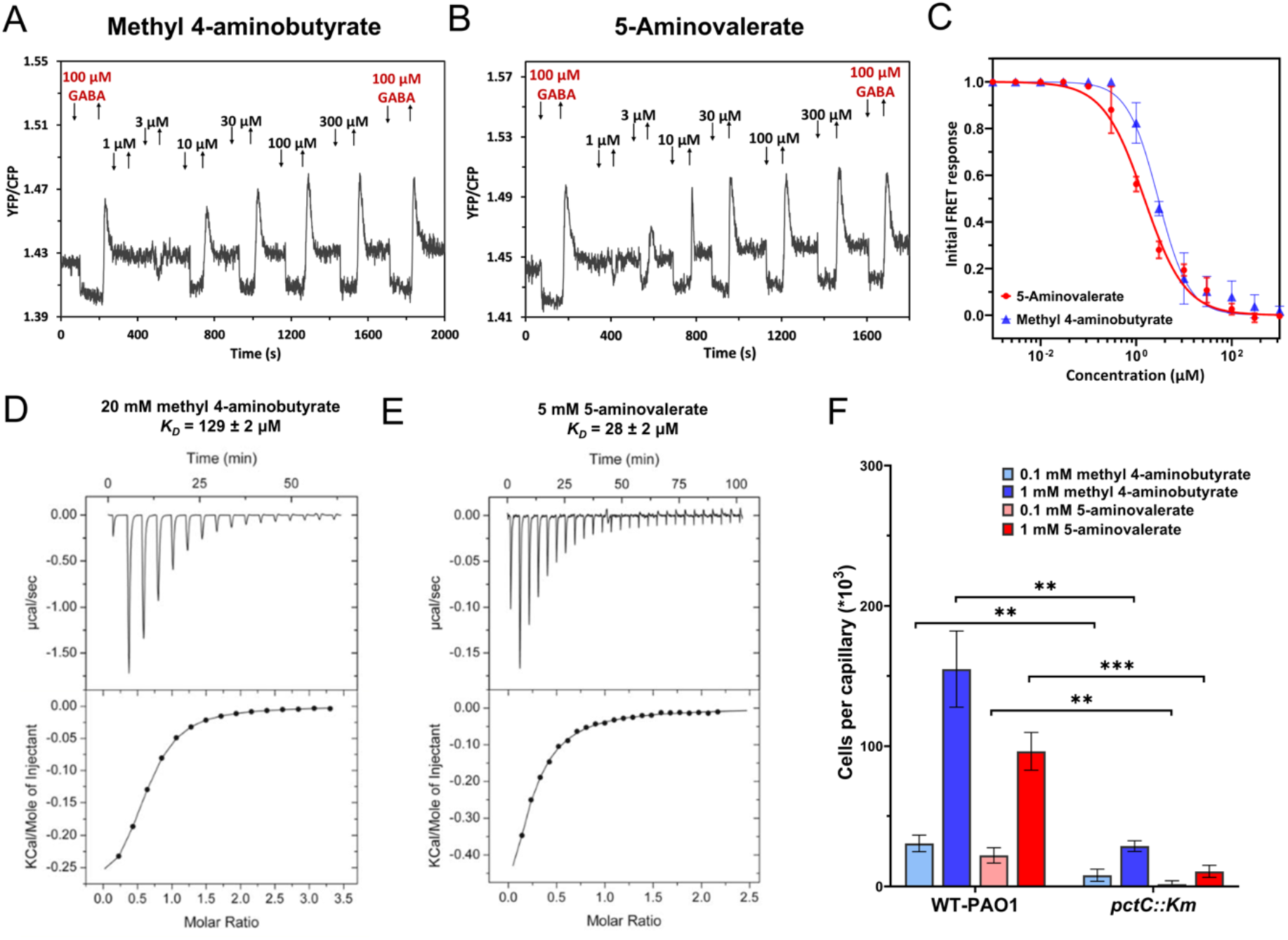
Characterization of PctC as a specific chemoreceptor for methyl 4-aminobutyrate and 5-aminovalerate. FRET measurements for *E. coli* cells expressing PctC-Tar hybrid as a sole receptor to the indicated concentrations of methyl 4-aminobutyrate (A) and 5-aminovalerate (B). The known ligand GABA (γ-Aminobutyric acid) was used as a positive ligand. (C) Corresponding FRET dose-response curves of PctC-Tar hybrid to methyl 4-aminobutyrate and 5-aminovalerate. Data were fitted using Hill equation, with the EC50 fit values being 2.9 ± 0.4 μM for methyl 4-aminobutyrate, and 1.5 ± 0.2 μM for 5-aminovalerate. The measurement and data analysis were conducted as described in the Fig. 4C. Microcalorimetric titrations of PctC-LBD with methyl 4-aminobutyrate (D) and 5-aminovalerate (E). The upper panel upper shows raw titration data, and lower shows integrated corrected peak areas of the titration data fit using the single-site binding model. (F) Capillary chemotaxis assays of the wildtype *P. aeruginosa* (WT-Hiroshima) and a *pctC* mutant strain (WT-Hiroshima was used as the parental strain) response to methyl 4-aminobutyrate and 5-aminovalerate. Data are shown as the means and standard deviations of results from three biological replicates each conducted in triplicate. Asterisks denote statistically significant differences; ns: non-significant, * p ≤ 0.05, ** p ≤ 0.01, *** p ≤ 0.001, **** p ≤ 0.0001.

### Characterization of PctP (PA1608) as a purine-specific chemoreceptor

The initial FRET screening showed that PA1608-Tar hybrid responded to guanine and inosine which are purine derivatives, whereas no responses were observed to cytosine, thymidine or uridine that all have a pyrimidine ring (Fig. 3A). Based on these observations, we speculated that PA1608 may be a purine-specific chemoreceptor. This hypothesis was further supported by TSA measurements of the binding of purine derivatives to PA1608, which all caused significant increases in the midpoint temperature of the protein unfolding transition (Fig. S3). We therefore further investigated the chemotactic responses of the PA1608-Tar hybrid to these 6 purine derivatives. When the PA1608-Tar hybrid was expressed as a sole chemoreceptor in *E. coli* FRET strain, clear attractant responses were observed to guanine and hypoxanthine in the lower micromolar range and to adenine at higher micromolar concentrations (Fig. 6A). Consistently, responses were seen for *E. coli* strain expressing PA1608-Tar in gradients of guanine and hypoxanthine, but not of adenine, using the microfluidic chemotaxis assay (Fig. 6B). FRET responses were also observed for PA1608-Tar expressing cells to all three corresponding purine nucleosides (Fig. 6C). No responses to these purine derivatives were observed for the *E. coli* FRET strain and microfluidic strain expressing only the wild type Tar receptor (Fig. S2D-I and Fig. S4). Overall, the extracellular domain of PA1608 is most sensitive to the two nucleobases guanine and hypoxanthine, followed by the purine nucleosides and finally by adenine (Fig. 6D).

**Fig. 6.**
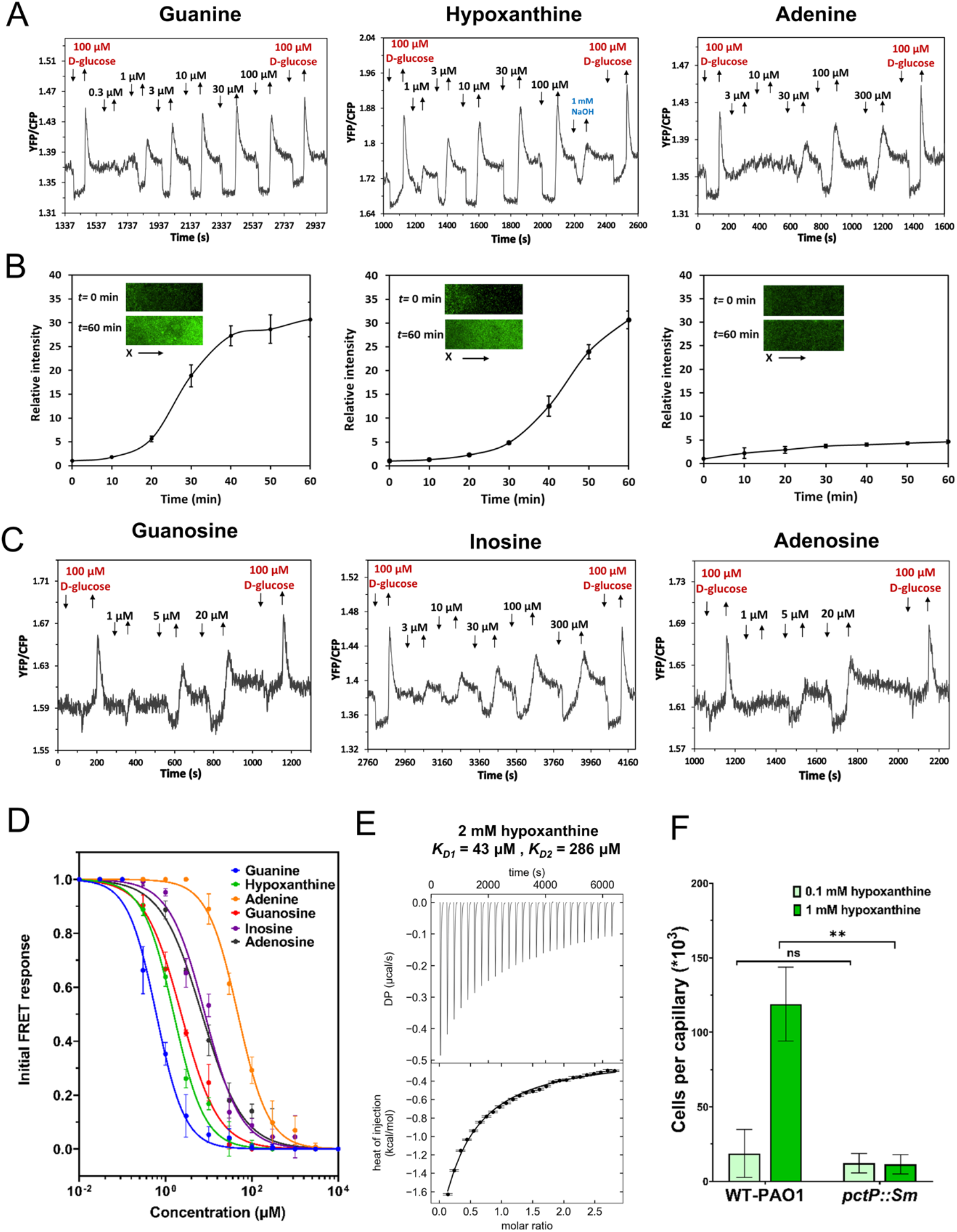
Characterization of PA1608 (PctP) as a purine-specific chemoreceptor. (A) FRET measurements for *E. coli* cells expressing PctP-Tar hybrid as a sole receptor to the indicated concentrations of guanine, hypoxanthine, and adenine. Since there is no previously characterized ligand for PctP, D-glucose was used as a positive control. The stock solutions of guanine and hypoxanthine were prepared with NaOH due to their poor solubility in neutral pH, and consequently 1 mM NaOH was used as control to exclude the influence of pH on the response of PctP-Tar hybrid. (B) Microfluidic assay of the chemotactic response of *E. coli* expressing PctP-Tar as the sole receptor and GFP as a label. Relative cell density (fluorescence intensity of GFP) in the observation channel over time in gradients of guanine, hypoxanthine, or adenine in the same order as in (A), with 50 mM in the source channel as indicated. Cell density in the observation channel before ligands stimulation (t=0) was used to normalize all data. Error bars indicate standard deviation from three independent biological replicates. Inserts show representative images of the observation channel at the beginning and the end of an experiment. The x-components (black arrow) indicate the direction up the concentration gradient. (C) FRET measurements as in (A) but for guanosine, inosine, and adenosine. (D) Dose-response curves for FRET measurements of responses mediated by PctP-Tar. Data were fitted using Hill equation, with the EC50 fit values being 0.6 ± 0.1 μM for guanine, 1.6 ± 0.2 μM for hypoxanthine, 47.6 ± 4.1 μM for adenine, 2.3 ± 0.3 μM for guanosine, 8.2 ± 1.9 μM for inosine, and 7.0 ± 1.3 μM for adenosine. The measurement and data analysis were conducted as described in Fig. 4C. (E) Microcalorimetric titrations of PctP-LBD with hypoxanthine. The upper panel shows the titration raw data, and the lower panel is the integrated dilution heat corrected and concentration normalized peak areas fitted with model for the binding with negative cooperativity to two symmetric sites. (F) Capillary chemotaxis assays of WT-PAO1 and a *pctP* mutant to 0.1 mM and 1 mM hypoxanthine. The *pctP* mutant was derived from the Washington parental strain that was used as a WT-PAO1 in this measurement. Data are shown as the means and standard deviations from three biological replicates each conducted in triplicate. Asterisks denote statistically significant differences; ns: non-significant, * p ≤ 0.05, ** p ≤ 0.01, *** p ≤ 0.001, **** p ≤ 0.0001.

Additional evidence for the specificity of PA1608 for purines was obtained by ITC, which showed that PA1608-LBD binds to hypoxanthine with high affinity in a process characterized by negative cooperativity (*K_D1_* = 43 μM and *K_D2_* = 286 μM) (Fig. 6E), while the binding of guanine and other purine derivatives was not detected due to the limited solubility or low binding affinity. Since hypoxanthine was not included in our initial screen to identify novel chemoeffectors, we have generated the PA1608 mutant and performed capillary assays of chemotaxis to hypoxanthine using the wild type and the mutant strains. Indeed, the wild type cells showed strong chemoattractant response to 1 mM hypoxanthine, whereas the inactivation of the PA1608 chemoreceptor nearly abolished this response (Fig. 6F). Taken together, PA1608 responded with different sensitivity to 6 purine derivatives and was thus renamed as PctP (*<u>P</u>seudomonas* <u>c</u>hemotaxis transducer for <u>p</u>urines).

### Characterization of chemoreceptors for ethanolamine, 2-phenylethylamine and other biogenic amines

Two remaining strong chemoattractants, ethanolamine (EA) and 2-phenylethylamine (PEA), did not elicit apparent responses in our initial FRET screening. In order to identify their specific chemoreceptor(s), we screened receptor mutant strains of *P. aeruginosa* in the presence of 20 mM EA and PEA using quantitative capillary assays. Although no conclusive results were obtained for EA, a significant decrease in the chemoattraction to PEA was observed for the *tlpQ* mutant strain, and intermediate reduction for PA4915, PA1646 and PA4520 mutant strains (Fig. 7A), indicating that these receptors might play a role in the chemotaxis to PEA. This was supported by the thermal shift assays, which suggested that TlpQ-LBD and PA4915-LBD might bind PEA, and PA4915-LBD might also bind EA (Fig. 7B). In these cases, we were unable to observe clear FRET responses of TlpQ-Tar and PA4915-Tar hybrids to PEA or EA, although these two hybrids showed good responses to D-glucose (Table S2) and TlpQ-Tar mediated (weak) attractant responses to its known ligands such as histamine, spermidine and spermine (Fig. S5A). This suggests that TlpQ-Tar and PA4915-Tar hybrids are not fully functional, explaining why responses to PEA and EA were not detected in our initial FRET screen. Nevertheless, our ITC measurements confirmed the direct binding of PEA to TlpQ-LBD and PA4915-LBD (Fig. 7C and D). Collectively, despite poor functionality of their hybrids, we conclude that TlpQ and PA4915 are the major chemoreceptors for PEA chemotaxis in *P. aeruginosa*.

**Fig. 7.**
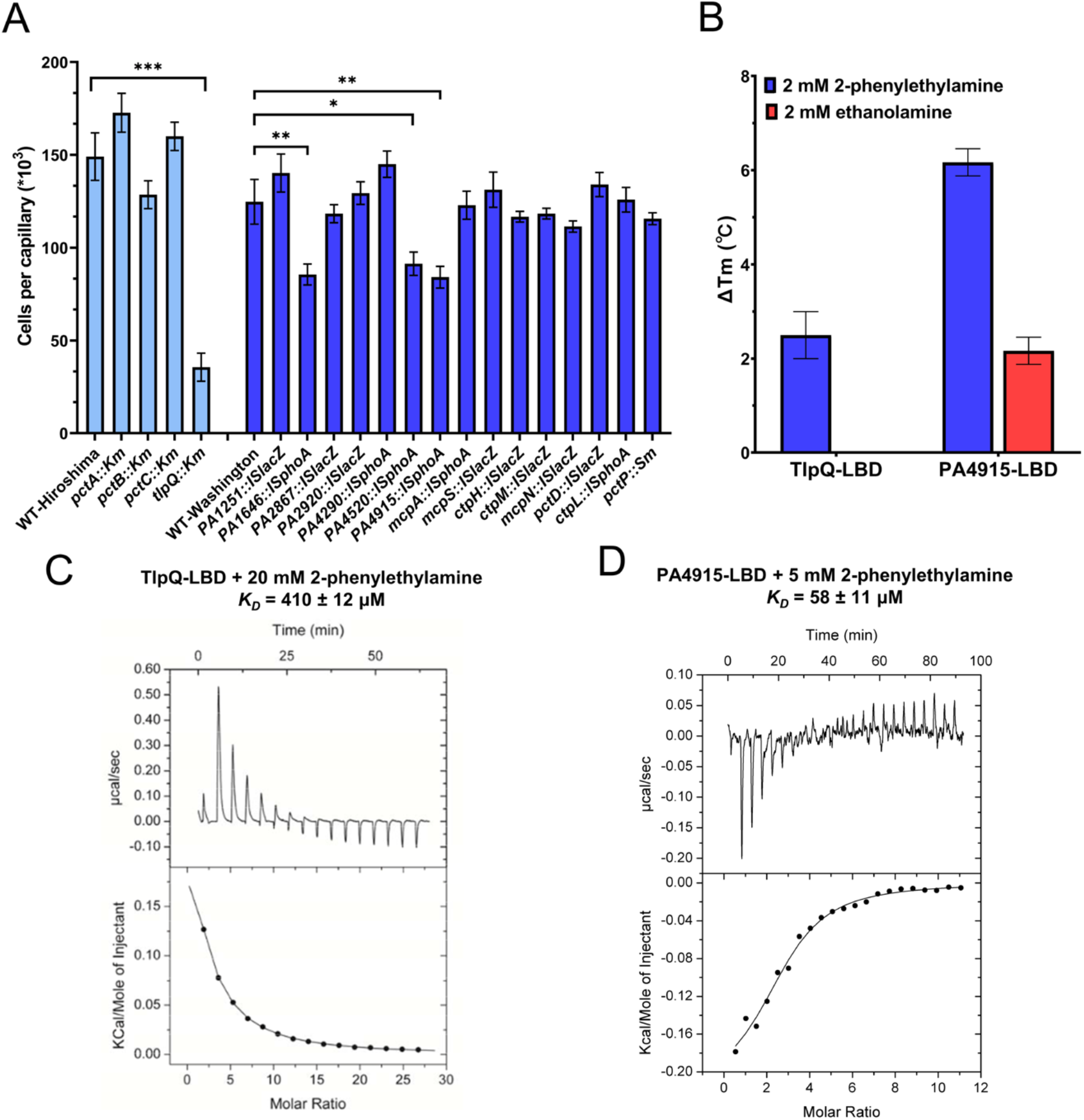
Identification of specific chemoreceptors for ethanolamine and 2-phenylethylamine. (A) Capillary assay for chemotaxis toward 20 mM 2-phenylethylamine (PEA) in different chemoreceptor mutants of PAO1. Mutant strains deficient in PctA, PctB, PctC and TlpQ were derived from WT-Hiroshima parental strain (shown in light blue) which was used as a reference. The remaining mutant strains were derived from WT-Washington parental strain (shown in blue) which was used as a reference for these mutants. Data are shown as the means and standard deviations of results from three biological replicates each conducted in triplicate. Asterisks denote statistically significant differences; ns: non-significant, * p ≤ 0.05, ** p ≤ 0.01, *** p ≤ 0.001, **** p ≤ 0.0001. (B) Thermal shift assay for TlpQ-LBD and PA4915-LBD in presence of 2 mM PEA or ethanolamine (EA). Data are shown as the means and standard deviations of results from three biological replicates each conducted in triplicate. (C, D) Microcalorimetric titrations of TlpQ-LBD (C) and PA4915-LBD (D) with PEA. Upper panels show raw titration data, whereas lower panels show the best fit of the integrated, concentration-normalized and dilution heat corrected raw data using the single-site binding model.

In previous studies, TlpQ was demonstrated to bind several biogenic amines, such as histamine, spermine, agmatine, cadaverine and putrescine (9). Since some of these compounds, as well as the newly identified ligand PEA, are produced by the decarboxylation of amino acids, we speculated that TlpQ might also sense tyramine, the decarboxylation product of tyrosine. Indeed, *tlpQ* mutant strain showed much reduced chemotaxis toward tyramine in the capillary assay (Fig. S5B), and binding of tyramine to TlpQ-LBD was confirmed by ITC (Fig. S5C).

Finally, we used FRET to test responses of amino acid chemoreceptors PctA, PctB and PctC to histamine, one of the most important biogenic amines (43, 44). A recent study (9) suggested that next to TlpQ, PctA and PctC also participate in the histamine chemotaxis, but microcalorimetric titrations of the corresponding LBDs revealed only binding to TlpQ. In contrast, FRET measurements confirmed that PctB and PctC can sense histamine in medium to high micromolar range (Fig. S6), highlighting the advantage of using receptor hybrids for characterization of low-affinity ligands.

## Discussion

Bacteria contain an extensive array of different sensory receptors that respond to a variety of stimuli, regulating multiple physiological functions including gene expression, chemotaxis or second messenger signaling. Major receptor families include sensory histidine kinases, chemoreceptors, adenylate, diadenylate and diguanylate cyclases and phosphodiesterases, as well as Ser/Thr/Tyr protein kinases and phosphoprotein phosphatases (45). Typically, these receptors are stimulated by the binding of signal molecules to their sensory domains that contain all the requisites for ligand binding. However, the lack of established signals that are recognized by receptors is currently a major bottleneck in our understanding of signal transduction processes in bacteria (46).

The fact that the same type of sensor domain is frequently found in different signal transduction systems suggests their modular nature and indicates that these domains have been exchanged among different receptor families during evolution. This notion is exemplified by the dCache domain that is the predominant type of the bacterial extracellular sensory domain, present in all major bacterial receptor families (47). Previous success with construction of hybrid receptors (35, 36, 48, 49) confirmed this modularity and indicated that the exchange of sensory domains between different receptors can also be reproduced in the laboratory, and it is likely that such hybrid construction could be extended beyond chemoreceptor-chemoreceptor and chemoreceptor-sensor kinase hybrids to sensor domains from other types of receptors. Ligand screening for hybrid receptors, as done in this work, has thus the promise to become a universal approach to identify receptor ligands and thus to tackle a major bottleneck in microbiology.

Here we combined the screening based on hybrid chemoreceptors with binding and capillary assays to systematically identify novel ligands of chemoreceptors in *P. aeruginosa*, an important pathogen with a complex lifestyle and the correspondingly broad chemosensory range. Since many known bacterial chemoattractants are metabolically valuable compounds (8, 37), we first performed high-throughput screenings for candidate chemoeffectors using the growth assay, followed by an evaluation of their potency as chemoattractants for *P. aeruginosa* in the capillary chemotaxis assays. The strongest chemoattractants were then prioritized for further investigation using a library of hybrid chemoreceptors, containing the sensor domains of *P. aeruginosa* chemoreceptors fused to the signaling domain of *E. coli* Tar, to identify the potential ligand-receptor pairs using FRET and microfluidic assays (35). This approach has several advantages, including highly sensitive and standardized chemotaxis assays already established in *E. coli*, which enable detection not only of high- but also of low-affinity ligands, and testing not only binding of ligands but also their signaling properties. It further avoids the complication of functional redundancy between *P. aeruginosa* chemoreceptors with overlapping ligand specificities. Nevertheless, this hybrid-based *in vivo* screening also has some limitations, primarily because most but not all of the constructed hybrids are functional, and in those cases it was complemented by *in vitro* ligand screening using TSA. Direct binding between the high-affinity ligands and sensor domains of chemoreceptors could further be confirmed using ITC. Finally, capillary chemotaxis assays were used to demonstrate the physiological relevance of identified ligand-receptor interactions for chemotaxis of *P. aeruginosa*.

This combination of assays enabled us to identify new ligands for previously studied chemoreceptors and to characterize new chemoreceptors in this well-studied model organism (Table 1). One example of the former category is the observed response to L-ornithine mediated by two amino acid chemoreceptors PctA and PctB, which support a recent study that implicated these receptors in *P. aeruginosa* chemotaxis towards L-ornithine (42). L-ornithine is a biologically versatile non-proteinogenic derivative of L-arginine. Besides its effects on growth, L-ornithine is known to promote of *P. aeruginosa* biofilm formation (50). L-ornithine might also accumulate in human lung environment during conditions that are associated with *P. aeruginosa* infections due to the elevated production of arginase (51), indicating a possible role of L-ornithine chemotaxis in virulence, and making its receptors potentially attractive target for therapeutic interventions. Indeed, the mutation of *pctA*, *pctB* and *pctC* reduced the accumulation of *P. aeruginosa* towards wounded lung epithelial cells (52). We observed that PctA senses L-ornithine at much lower concentrations than PctB, which is consistent with the previously observed difference in the sensitivity of these two receptors to L-arginine and several other amino acids (53). Another example is the *P. aeruginosa* response to other amino acid derivatives, methyl 4-aminobutyrate and 5-aminovalerate. We could show that these compounds are sensed by PctC, the high-affinity receptor for gamma-aminobutyric acid (GABA). This is consistent with the structural similarity of these compounds to GABA, although their affinities to PctC are 10 to 100-fold lower than that of GABA (54).

Besides assigning new ligands to known chemoreceptors, we identified a novel purine-specific chemoreceptor PctP (PA1608). FRET results show that this receptor has highest affinity for guanine and hypoxanthine, intermediate affinity for purine nucleosides and significantly lower affinity for adenine, and it exhibits no response to pyrimidine derivatives. A lacking availability of nucleotide bases might limit growth of bacterial pathogens in the human host (55), including growth of *P. aeruginosa* in the lungs of cystic fibrosis (CF) patients (56), and cross-feeding of purine derivatives might play a role in a polymicrobial community in CF lungs (57). Thus, purine derivatives are both important nutrients and might be available during *P. aeruginosa* infection. Despite their potential importance, there are only few existing reports of chemotaxis towards purine or pyrimidine derivatives (11, 12, 58, 59), and only a single bacterial chemoreceptor specific for metabolizable purines has been characterized so far, McpH from *Pseudomonas putida* KT2440 (59). Interestingly, despite the phylogenetic proximity of *P. aeruginosa* and *P. putida*, McpH possesses a dCache-type LBD, whereas PctP has a four-helix bundle type LBD, indicative of convergent evolution (60) and providing further support to the important role of purine chemotaxis.

Finally, *P. aeruginosa* response to ethanolamine, 2-phenylethylamine and tyramine expands the ligand range of characterized chemoreceptors. TlpQ was previously reported to bind several biogenic amines (BAs), including putrescine, histamine, agmatine and cadaverine (9), which are derived from the decarboxylation of L-amino acids. BAs have important physiological functions in eukaryotic and prokaryotic cells, and many bacteria are able to produce and/or degrade Bas (44, 61). Here, with PEA and tyramine we identified two additional TlpQ ligands. Sequence comparison between the dCache domain of TlpQ and the amino acid-binding dCache domains of PctA, PctB and PctC (62) shows conservation of amino acids that bind the amine group, which may explain the capacity of these receptor to bind either amino acids or their decarboxylated derivatives. Furthermore, we identified a second, previously uncharacterized, receptor PA4915 as a sensor of PEA, and possibly also of EA. The binding affinity of PA4915-LBD for PEA is even higher than that of TlpQ LBD, suggesting that it might be particularly important to mediate response to low concentrations of PEA. Notably, PA4915 possesses the four-helix bundle type LBD, which provides an interesting example of receptors within one bacterium that harbor different types of LBD but respond to the same ligand.

In addition to characterizing novel receptor specificities for chemical ligands, we observed that multiple LBDs of *P. aeruginosa* chemoreceptors can mediate specific responses to pH. In two model neutrophilic bacteria where pH taxis has been studied, *E. coli* (20) and *Bacillus subtilis* (19), bacterial accumulation toward neutral pH is ensured by opposite pH responses mediated by different receptors. In *E. coli*, the acidophilic response is primarily mediated by the LBD of Tar and the alkaliphilic response by the LBD of Tsr (20). Our results for the hybrid receptors suggest the existence of similar, but possibly more complex, bidirectional pH taxis in *P. aeruginosa*, with the LBDs of PctC and PA1646 mediating acidophilic taxis and those of PctB and PctP mediating alkaliphilic taxis. Particularly interesting are pH responses of the hybrids containing LBDs of PctA and TlpQ, which show repellent responses to both high and low pH, indicating that these receptors might even be individually able to mediate bacterial accumulation toward neutral pH. Given this multitude of pH responses mediated by the LBDs of *P. aeruginosa* chemoreceptors, along with recently described general effects of pH on chemoreceptor LBDs (18), it would be interesting to investigate behavior of *P. aeruginosa* and other bacteria with complex chemosensory systems in pH gradients in future studies.

Taken together, our systematic screening strategy enabled us to enlarge the spectrum of ligand specificities of sensor domains in *P. aeruginosa*, despite it being already well-studied model for bacterial chemotaxis. A similar strategy could be applied for the systematic characterization of unknown sensor domains from different types of receptors in other species, including those that are unculturable.

## Materials and methods

### Bacterial strains, plasmids and culture conditions

Bacterial strains and plasmids are listed in Table S3. For chemotaxis and FRET experiments, *E. coli* was grown in TB medium (1% tryptone and 0.5% NaCl) at 34 °C. For molecular cloning and protein expression, *E. coli* was grown in LB medium at 37 °C. *P. aeruginosa* was grown overnight in M9 minimal medium containing 15 mM D-glucose at 37 °C. When necessary, antibiotics were used at the following final concentrations: kanamycin, 50 µg/ml (*E. coli* strains); ampicillin, 100 µg/ml (*E. coli* strains); chloramphenicol, 34 µg/ml (*E. coli* strains) and 100 µg/ml (*P. aeruginosa* strains); tetracycline, 50 µg/ml (*P. aeruginosa* strains).

### PA1608 mutant generation

To generate the *pctP* mutant strain, a 1280-bp fragment of PA1608 was amplified by PCR using primers PA1608_MUT_F and PA1608_MUT_R (Table S3). The PCR product was cloned into pGEM^®^-T vector and transformed into *E. coli* DH5α. The resulting plasmid pGEMT®-PA1608 was digested with ApaI and SpeI and the insert cloned into pKNG101 previously digested with the same enzyme. The resulting plasmid pKNG101-PA1608 was then transformed into *E. coli* CC118 λpir. pKNG101_PA1608 was then introduced into *P. aeruginosa* PAO1 by electroporation according to (63). The mutant was verified by PCR and sequencing.

### Growth assays

The analysis of the nutritional profile was carried out using the “Phenotype Microarrays^TM^” plates PM1, PM2A and PM3B (for further information, refer to https://www.biolog.com/wp-content/uploads/2020/04/00A-042-Rev-C-Phenotype-MicroArrays-1-10-Plate-Maps.pdf). Each of these plates contains 95 chemical compounds and one control (H_2_O). To determine the growth of *P. aeruginosa* using these compounds as the sole carbon or nitrogen source, the lyophilized compounds present on the Biolog plates were resuspended in 90 µL of either M9 medium (for Biolog PM1 and PM2A plates) or M8 medium (M9 minimal medium without NH_4_Cl) containing 15 mM glucose (for Biolog PM3B plate). Subsequently, a *P. aeruginosa* overnight culture was washed twice with M8 salts medium, diluted to an OD_600_ of 0.2 in M9 or M8 salts medium and then the wells with each of the compounds were inoculated with 10 µL of these cultures. Finally, growth was monitored at 37 °C with shaking, determining the OD_600_ every hour for 48 h in a Bioscreen Microbiological Growth Analyser (Oy Growth Curves Ab Ltd, Helsinki, Finland).

### Chemotaxis capillary assays for P. aeruginosa strains

Overnight cultures in M9 minimal medium supplemented with 6 mg/L Fe-citrate, trace elements (64), and 15 mM glucose were used to inoculate fresh medium to an OD_660_ of 0.05. Cells were cultured at 37 °C to an OD_660_ of 0.4. Subsequently, cells were washed twice by centrifugation (1,667 × g for 5 min) and resuspended in chemotaxis buffer (50 mM KH_2_PO_4_/K_2_HPO_4_, 20 mM EDTA, 0.05% [vol/vol] glycerol, pH 7.0). Aliquots (230 μl) of the cell suspension at an OD_660_ of 0.1. were placed into the wells of 96-well microtiter plates. Then, 1-μl capillaries (Microcaps, Drummond Scientific) were heat-sealed at one end and filled with buffer (control) or chemoeffector solution prepared in chemotaxis buffer. The capillaries were rinsed with sterile water and immersed into the bacterial suspensions at their open ends. After 30 min, capillaries were removed from the wells, rinsed with sterile water, and emptied into 1 ml of chemotaxis buffer. Serial dilutions were plated onto M9 minimal medium plates supplemented with 20 mM glucose, incubated at 37 °C prior to colony counting. Data were corrected with the number of cells that swam into buffer containing capillaries.

### Construction of hybrid chemoreceptors

To construct the hybrid type 1 in a high-efficient way, the drop-out plasmid pHC8 was generated by Golden Gate Assembly kit (New England BioLabs), and the basic genetic parts are from Marburg collection (65). Dropout part served as placeholder, which carried a full expression cassette for the fluorescent proteins sfGFP to enable visible distinction of correct colonies and outward facing BsaI recognition sites. The signaling domain of Tar [198–553] was connected after the dropout part. The extracellular sensory domains of *P. aeruginosa* chemoreceptors containing BsaI recognition sites at both ends were synthesized, which allowed for replacing the dropout part by Golden Gate reaction, then generating the hybrid chemoreceptor in one-step reaction. For the constructions of hybrid type 2 and type 3, the coding sequences of each hybrid were amplified using PCR reaction (oligonucleotide sequences are shown in Table S3). The amplified fragments containing overlapping sequences of vector pKG116 were ligated into the digested vector pKG116 using Gibson assembly reaction in NEBuilder® HiFi DNA Assembly Master Mix (New England BioLabs). After cloning, the active or functional hybrid type 3 were selected from a library of chemoreceptor[1–X]-XXXXX-[203– 553] as described previously (35).

### FRET measurements

FRET measurements were performed as described previously (35, 38, 66%). Cultures of the receptorless *E. coli* strain VS181 expressing chimeras of interest and CheY-YFP/CheZ-CFP FRET pair were prepared by inoculating 200 μl of the overnight culture into 10 ml TB medium supplemented with appropriate antibiotics and inducers (50 µM isopropyl-β-D-thiogalactoside (IPTG) and 1-2 µM sodium salicylate), and grown in a rotary shaker at 34°C and 275 rpm. Cells were harvested at OD_600_ of 0.5 by centrifugation, and washed twice with tethering buffer (10 mM KH_2_PO_4_/K_2_HPO_4_, 0.1 mM EDTA, 1 μM methionine, 10 mM sodium lactate, pH 7.0). For microscopy, the cells were attached to the poly-lysine-coated coverslips for 10 min and mounted into a flow chamber that was maintained under constant flow of 0.3 ml/min of tethering buffer using a syringe pump (Harvard Apparatus) that was also used to add or remove compounds of interest. The pH value of all tested compounds was adjusted to 7, except for compounds that are only soluble under acidic or basic conditions, where the background buffer at corresponding pH was tested as a negative control. FRET measurements were performed on an upright fluorescence microscope (Zeiss AxioImager.Z1) equipped with photon counters (Hamamatsu). The fluorescence signals were recorded and analyzed as described previously (17).

### Protein overexpression and purification

The LBDs of chemoreceptors PctA, PctB, PctC, PA4915 and TlpQ were purified as described previously (9, 67). For the remaining proteins, the LBDs were cloned into a pET28b(+) expression vector. *E. coli* BL21 (DE3) harboring the LBD expression plasmid was grown in 5 L Erlenmeyer flasks containing 1 L LB medium supplemented with kanamycin under continuous stirring (200 rpm) at 37 °C. When OD_600_ reached 0.6, 0.1 mM isopropyl-β-D-thiogalactoside (IPTG) was added to induce protein expression. Growth was continued at 16 °C for 12 h and cells were harvested by centrifugation at 10 000 x *g* for 30 min at 4 °C. Proteins were purified by metal affinity chromatography using modified procedures for His GraviTrap^TM^ column (Cytiva lifesciences, Marlborough, Massachusetts, USA). Briefly, cell pellets were resuspended in binding buffer (20 mM sodium phosphate, 500 mM NaCl, and 20 mM imidazole, pH 7.4) supplemented with 0.2 µg/ml lysozyme, 1 mM MgCl_2_, 1 mM PMSF, stirred for 30 min at 4 °C and lysed by an ultrasonic homogenizer for 10 min at 80 % amplitude (SONOPULS HD 4000; BANDELIN electronic GmbH & Co. KG, Berlin, Germany). After centrifugation at 20 000 x *g* for 30 min, the supernatant was loaded into His GraviTrap^TM^ column pre-equilibrated with binding buffer, and target proteins were eluted by elution buffer (20 mM sodium phosphate, 500 mM NaCl, and 500 mM imidazole, pH 7.4). The protein fractions were dialyzed with the dialysis buffer (10 mM sodium phosphate, pH 5.5) to remove imidazole and NaCl, and concentrated using Amicon Ultra-15 centrifugal filters (Merck Millipore Ltd, Burlington, Massachusetts, USA).

### Soft agar plate gradient assay for E. coli strains

To establish gradients, 200 µl aliquots of 100 mM chemical solutions were applied to the center line of minimal A agar plates (0.25% (w/v) agar, 10 mM KH_2_PO_4_/K_2_HPO_4_, 8 mM (NH4)_2_SO_4_, 2 mM citrate, 1 mM MgSO_4_, 0.1 mg/ml of thiamine-HCl, 1 mM glycerol, and 40 μg/ml of a mixture of threonine, methionine, leucine, and histidine) supplemented with antibiotics and inducers and incubated overnight at 4 °C for gradient formation. The receptorless *E. coli* cells expressing the chimera of interest as a sole receptor were grown overnight in 5 ml TB medium, harvested by centrifugation, washed by tethering buffer, resuspended in 200 µl tethering buffer, and applied to the plate at a 2.5 cm distance from the line where the chemical was inoculated. Plates were incubated at 30 °C for 24−48 hours.

The screening of active or functional hybrid type 3 containing a random linker was performed as described previously (35). The library of hybrids mixture was applied to an agar plate with D-glucose gradients for the three rounds of selection. Cells that accumulated at the edge of the colony ring were selected and re-inoculated for the second round of selection. After three rounds of selection, the best chemotactic cells were streaked out on LB plates to obtain single colonies. The chemotaxis behavior of the selected colonies was further confirmed on soft agar plates. The sequence of random linker for the functional hybrid was identified by DNA sequencing.

### Microfluidic assays

The microfluidic assay was performed as previously described, using a chip with 24 parallel microchannels (68). The receptorless *E. coli* strain UU1250 expressing GFP and chimera of interest were grown at 34 °C in TB supplemented with antibiotics and inducers until OD_600_ of 0.5. Cells were harvested by centrifugation and washed twice with tethering buffer. The compounds of interest were dissolved in tethering buffer at a concentration of 50 mM and the pH was adjusted to 7.0. The chemical source microchannels were filled with 4 % (w/v) low-gelling temperature agarose to create a semi-permeable barrier. *E. coli* cells were added in the reservoir well and allowed to spread for 30 min into the channels. Compounds were added to the source well and allowed to form a concentration gradient. Cell fluorescence was recorded with a Nikon Ti-E inverted microscope system (Nikon Instruments Europe BV, Amsterdam, Netherlands) using a 20x objective. Data were analyzed using ImageJ (Wayne Rasband, National Institutes of Health, USA).

### Thermal shift assays

Thermal shift assays were performed in 384 microtiter plates using a Bio-Rad CFX384 Touch^TM^ Real-Time PCR instrument with the presence or absence of potential chemoattractants. Each 25 μl assay mixture contained 20.5 μl purified protein (30-100 μM) in phosphate buffer (10 mM sodium phosphate, pH 5.5), 2 μl SYPRO^TM^ orange (Invitrogen by Thermo Fisher Scientific, Waltham, Massachusetts, USA) at 5 × concentration, and 2.5 μl 20 mM potential chemoattractant. Samples were heated from 23 °C to 95 °C at a scan rate of 1 °C/min. The protein unfolding curves were monitored by detecting changes in SYPRO^TM^ Orange fluorescence. The resulting data permitted the calculation of the mid-point of the protein unfolding transition, or melting temperatures (Tm) using the first derivative values from the raw fluorescence data, which was analyzed by the Bio-Rad CFX manager 3.1 software.

### Isothermal titration calorimetry

Experiments were conducted on a VP-microcalorimeter (Microcal, Amherst, MA, USA) at the temperatures indicated in Table S4. Proteins were dialyzed into the buffer specified in Table S4 and placed into the sample cell. Ligand solutions were made up in dialysis buffer at the concentrations indicated in Table S4 and titrated into the protein. The mean enthalpies measured from the injection of ligands into buffer were subtracted from raw titration data prior to data analysis with the MicroCal version of ORIGIN. Data were fitted with the single-site binding model. In cases were data analysis with this model did not result in a satisfactory fit, data were analyzed in SEDPHAT (69) using a model for the binding with negative cooperativity to a macromolecule containing two symmetric sites.

## Data availability

All of the data are included in this article.

## Acknowledgements

This work was supported by the Max Planck Society and the Hessian Ministry of Higher Education, Research and the Arts (HMWK) – LOEWE research cluster “Diffusible Signals,” subproject A1 (to VS), the Spanish Ministry for Science and Innovation/*Agencia Estatal de Investigación* 10.13039/501100011033 (grants PID2020-112612GB-I00 to TK and PID2019-103972GA-I00 to MAM) and the Junta de Andalucía (grant P18-FR-1621 to TK). JPCV was supported by the grant Unión Europea-NextGenerationEU RD 289/2021 UPM-Recualifica Margarita Salas. WX was supported by Peterson Group “Serving Hometown” Elites scholarship of Peterson Group Charity Foundation Limited and Chinese Scholarship Council (CSC) scholarship.

## Abbreviations

FRET: Förster resonance energy transfer
ITC: isothermal titration calorimetry
LBD: ligand binding domain
TSA: thermal shift assays
PEA: 2-phenylethylamine
EA: ethanolamine
BAs: biogenic amines.

## Conflict of interest

The authors do not declare any conflict of interest.

## Supplementary Data

**Fig. S1.**
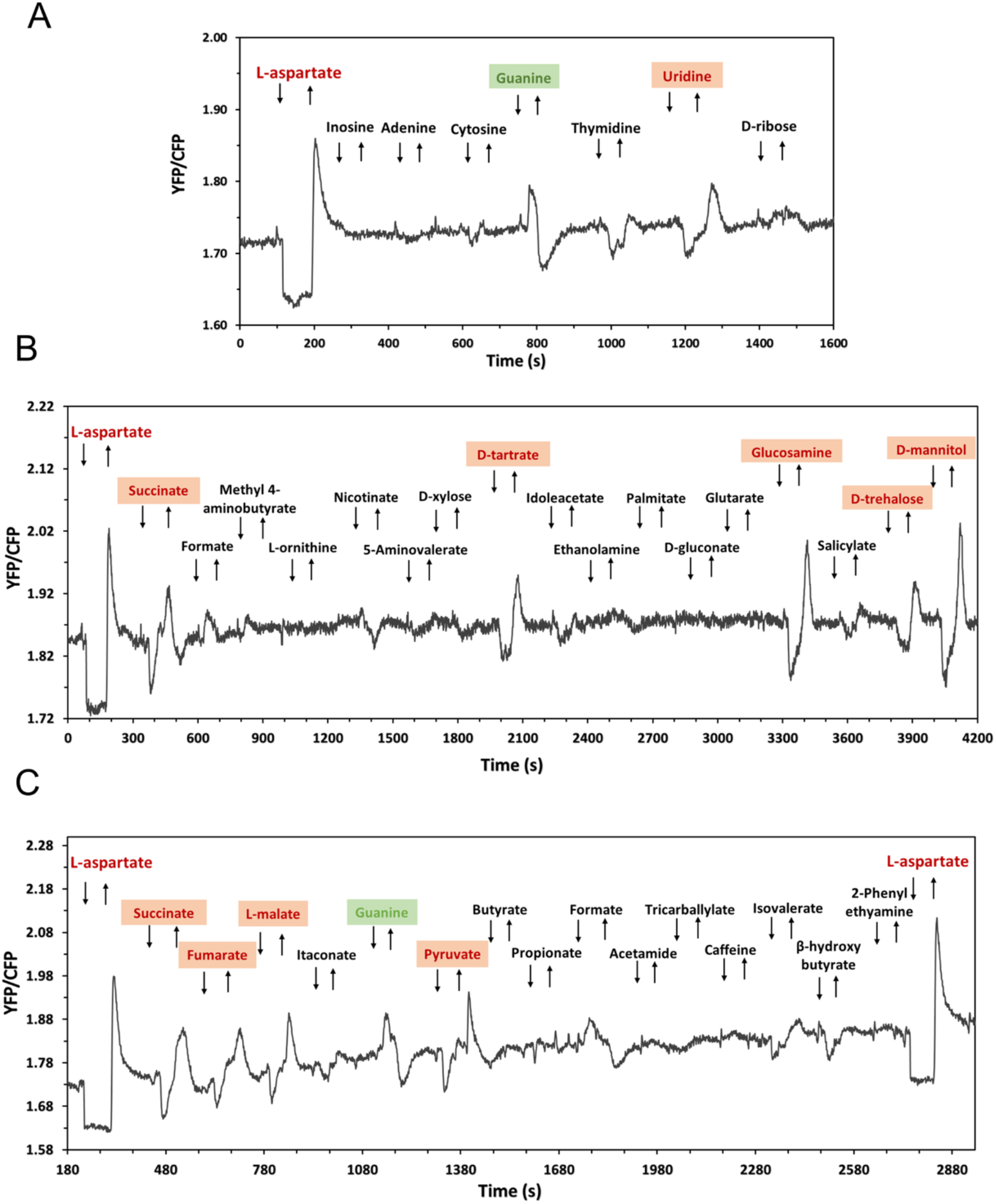
Responses of Tar control to unassigned potential ligands. FRET measurements of responses in receptorless *E. coli* cells expressing wildtype Tar as the sole receptor, stimulated by the stepwise addition (down arrow) and subsequent removal (up arrow) of 100 µM L-aspartate (L-Asp, positive ligand for Tar) or other tested compounds, as indicated. The chemoattractants are shown in red and the chemorepellents are shown in green.

**Fig. S2.**
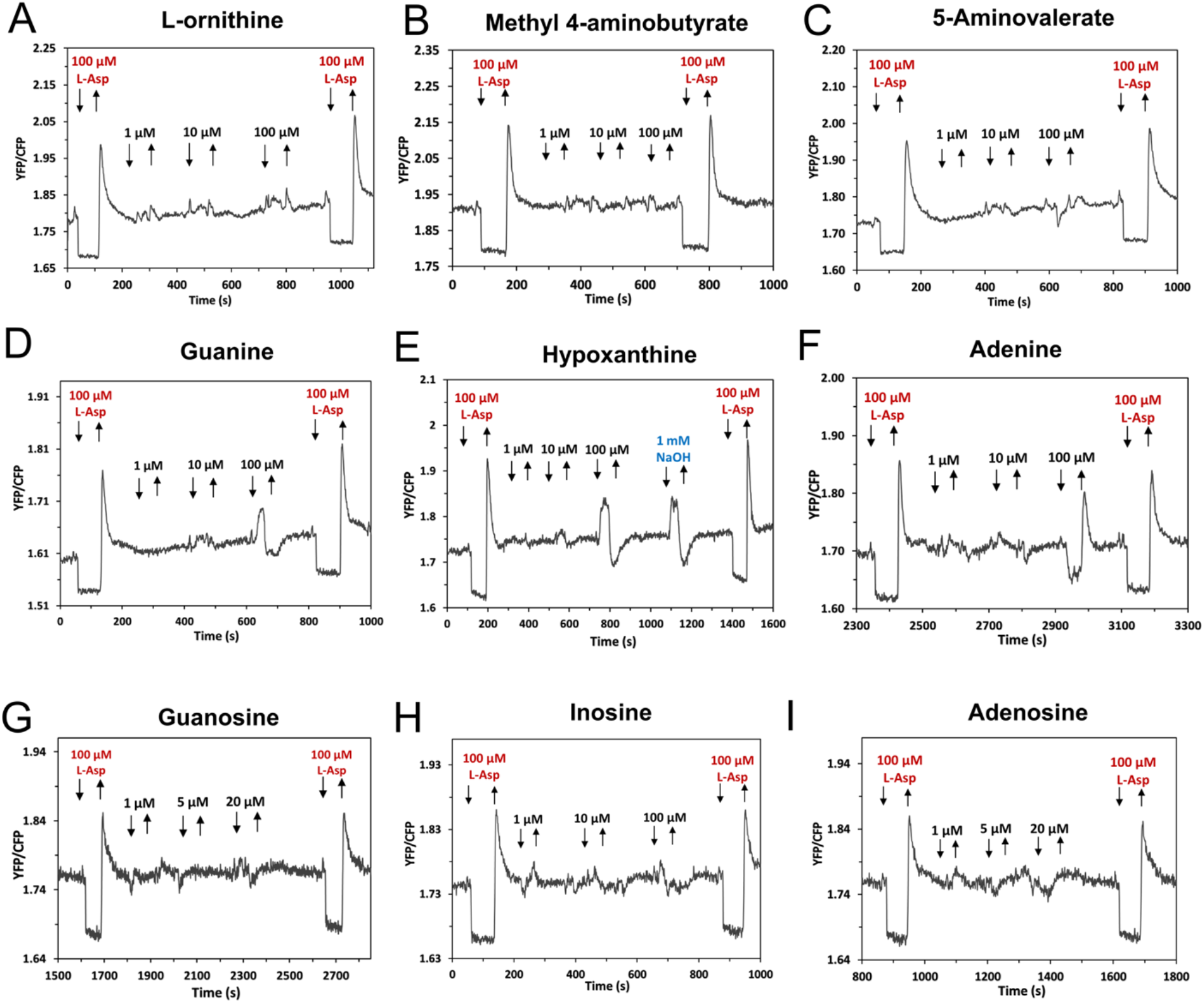
Responses of Tar control to different doses of characterized ligands. FRET measurements of responses in receptorless *E. coli* expressing Tar as a sole receptor to indicated concentrations of different compounds. 100 µM L-aspartate (L-Asp) were used as positive ligand for Tar. Since guanine and hypoxanthine are only soluble at high concentrations in alkalic solutions, an equivalent concentration of NaOH was used as negative control.

**Fig. S3.**
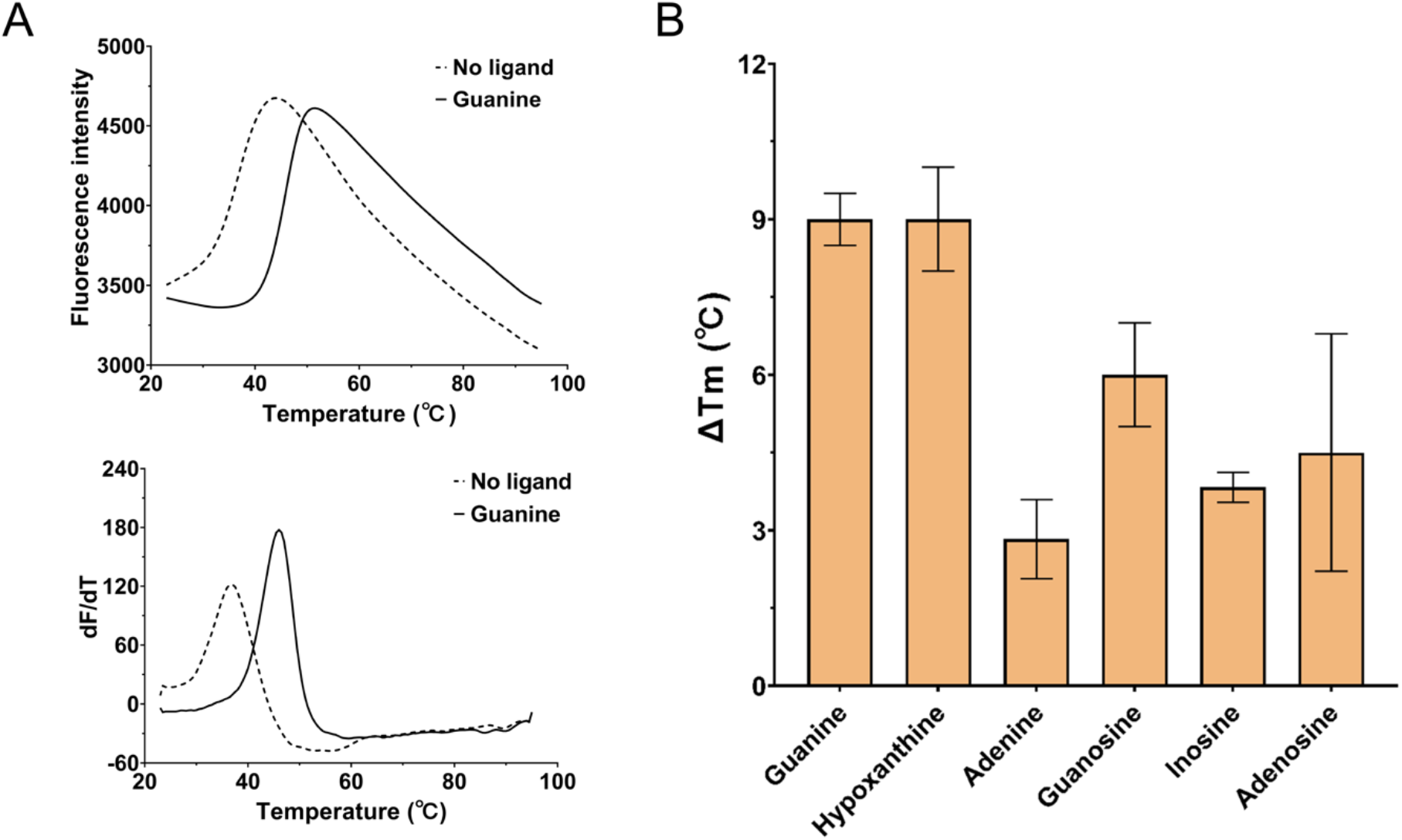
Thermal stability of PctP-LBD is enhanced by purine derivatives. Thermal shift assay for purified PctP-LBD in presence of 2 mM guanine, hypoxanthine, adenine, guanosine, inosine or adenosine. (A) The thermal unfolding curves of PctP-LBD in the absence and presence of guanine; upper panel: raw data, lower panel: first derivative of raw data. (B) Increases in the midpoint of protein unfolding transition (Tm) induced by different ligands. Data shown are the means and standard deviations from three biological replicates conducted in triplicate.

**Fig. S4.**
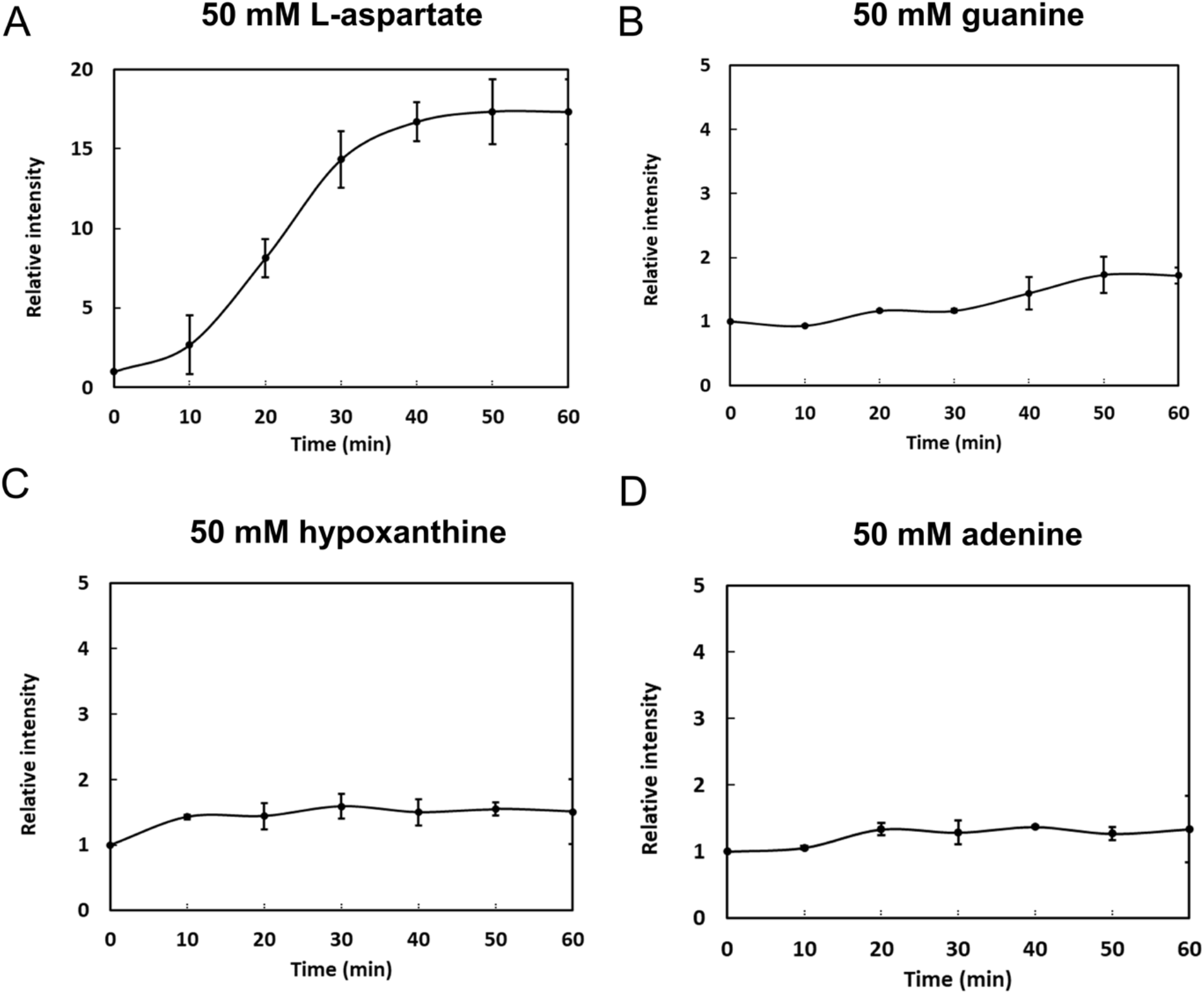
Microfluidic assays of the chemotactic responses of *E. coli* expressing Tar as the sole receptor. Relative cell density (fluorescence intensity) in the observation channel over time in gradients of L-aspartate, guanine, hypoxanthine, and adenine with 50 mM in the source channel.

**Fig. S5.**
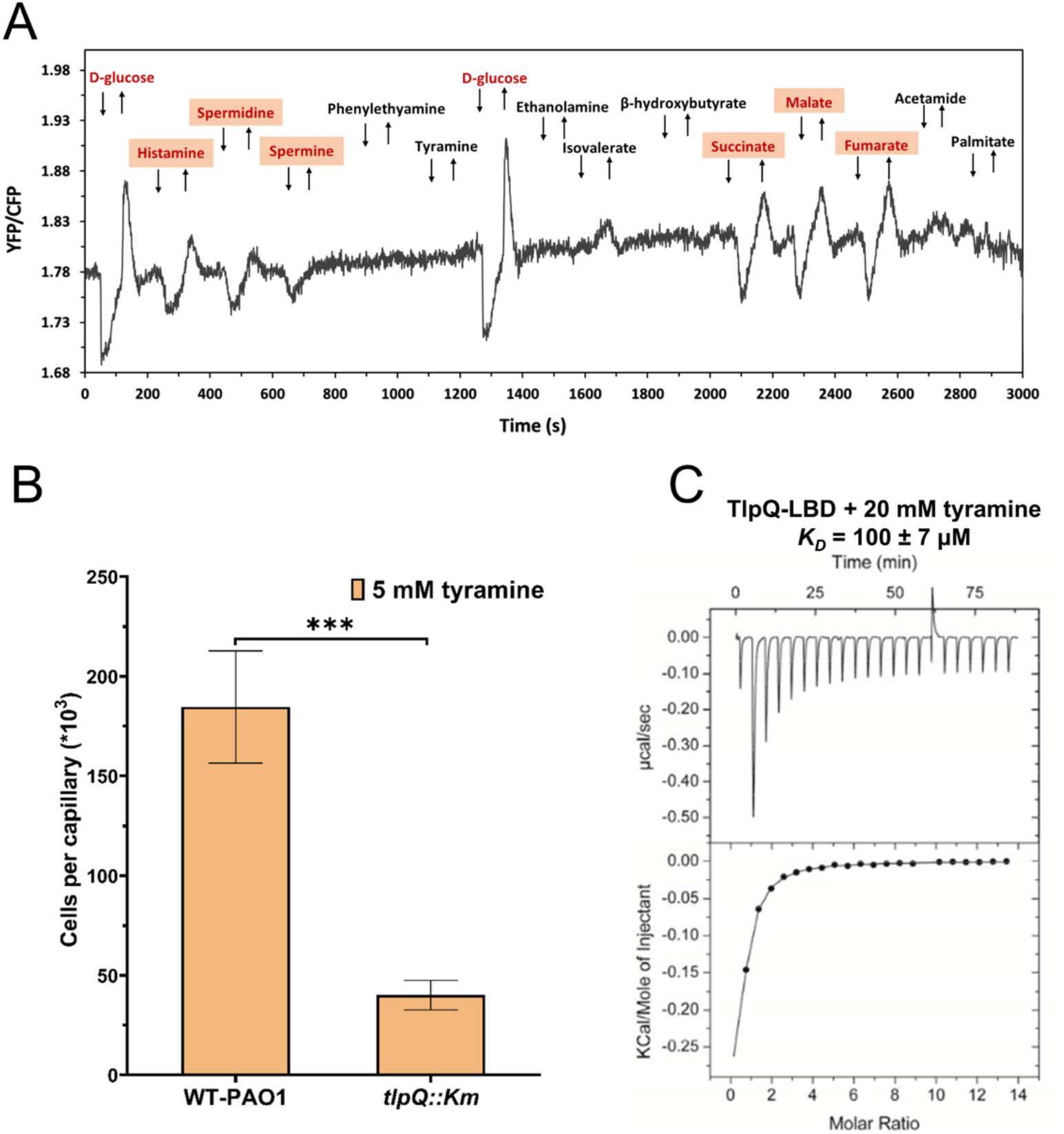
Characterization of TlpQ as a specific chemoreceptor for tyramine. (A) Screening of unassigned potential ligands for TlpQ-Tar using FRET. FRET measurement was performed with receptorless *E. coli* cells expressing TlpQ-Tar as the sole receptor exposed to a stepwise addition (down arrow) and subsequent removal (up arrow) of 100 µM D-glucose (positive ligand for chimeras), three known ligands (histamine, spermine and spermidine) and other chemical compounds. The chemoattractants were shown in red. (B) Capillary chemotaxis assays of *tlpQ* deficient strain and WT-PAO1 (WT-Hiroshima) to 5 mM tyramine. Data are shown as the means and standard deviations from three biological replicates conducted in triplicate. (C) Microcalorimetric studies showing the binding of tyramine to TlpQ-LBD. The lower panels are the integrated, dilution heat corrected and concentration normalized peak areas fitted with the single-site binding model.

**Fig. S6.**
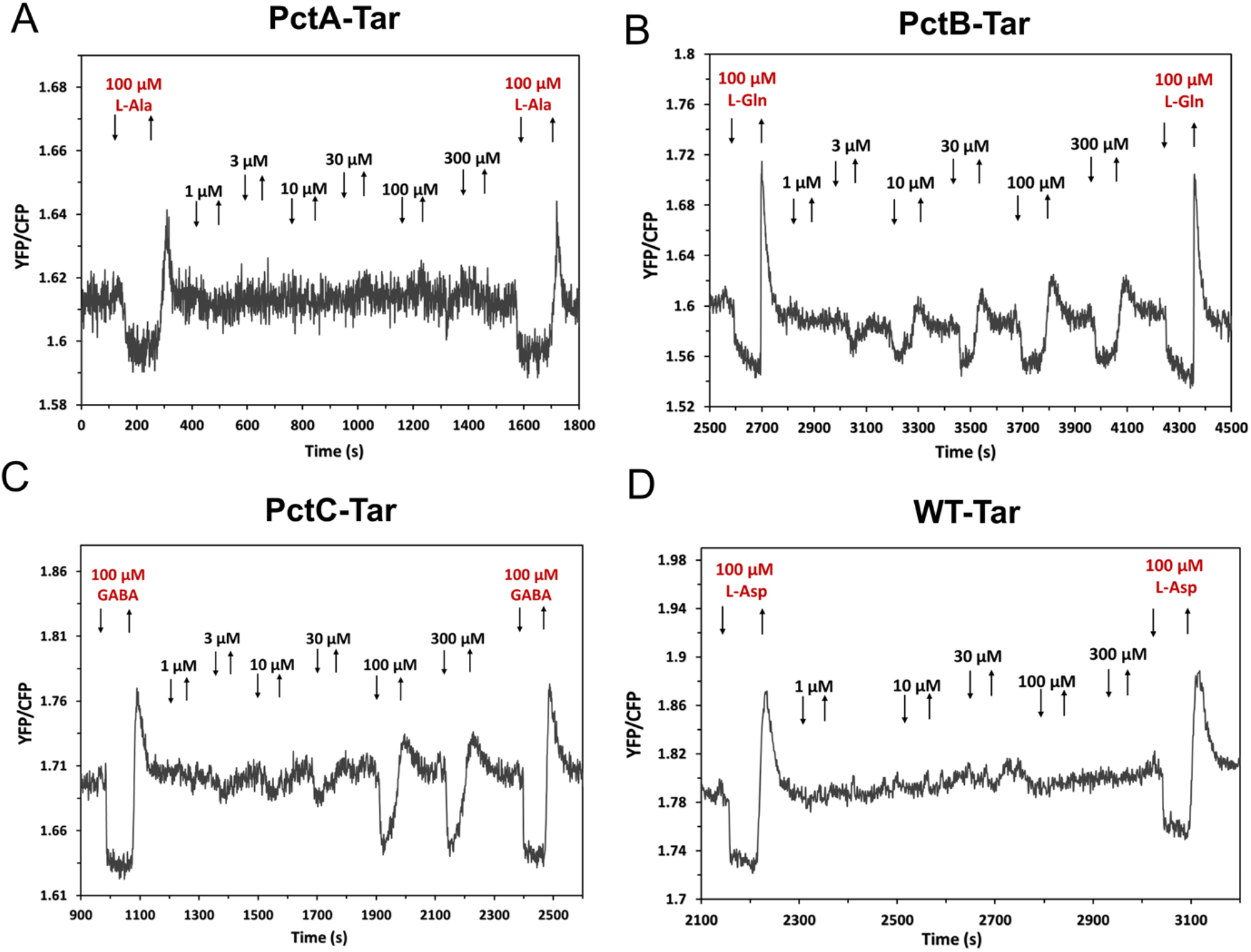
PctB and PctC mediate responses to histamine. FRET measurements of responses mediated by PctA-Tar (A), PctB-Tar (B), PctC-Tar (C) or Tar (D) as a sole receptor to indicated concentrations of histamine. 100 µM L-aspartate (L-Asp) were used as positive control for Tar.

**Table S1 Growth of P. *aeruginosa* PAO1 in M9 minimal medium supplemented with each of the compounds present in the Biolog plates PM1, PM2A and PM3B as nitrogen source (in nitrogen-free medium) or carbon source (in carbon-free medium).** There are 202 that supported bacterial growth at different levels. Compounds were divided into different groups according to the magnitude of growth. The 39 highlighted compounds were further studied using quantitative capillary chemotaxis assays (Fig. 1).

**Table S2 Summary of FRET responses of 16 hybrid chemoreceptors in the presence of 100 μM ligands.** “A”: attractant response; “R”: repellent response; “*”: strong response; “*<u>R or A</u>”*: weak response (or unspecific response); “--”: no response; “blank”: no analysis. The yellow highlights were considered as positive ligands.

**Table S4. Experimental conditions used for microcalorimetric titrations.**

**Table S3 Strains, plasmids and oligonucleotides used in this study.**

## Notes

### Competing Interest Statement

The authors have declared no competing interest.

### Summary of Updates

Supporting information added; authors list corrected

