## Supplemental Table 1 for "Systematic mapping of chemoreceptor specificities for *Pseudomonas aeruginosa*"

**Table S1 Growth of *P. aeruginosa* PAO1 in M9 minimal medium supplemented with each of the compounds present in the Biolog plates PM1, PM2 and PM3 as nitrogen source (in nitrogen-free medium) or carbon source (in carbon-free medium).** There are 202 that supported bacterial growth at different levels. Compounds were divided into different groups according to the magnitude of growth. The 39 highlighted compounds were further studied using quantitative capillary chemotaxis assays (Fig.1).

| **No.** | **High bacterial growth (OD_600nm_>0.25)** | **Medium bacterial growth (0.25>OD_600nm_>0.1)** | **Low bacterial growth (0.1<OD_600nm_<0.05)** |
| --- | --- | --- | --- |
| 1 | p-Hydroxy Phenyl Acetic Acid | Citric Acid | Mucic Acid |
| 2 | D, L-Malic Acid | L-Malic Acid | **Tricarballylic Acid** |
| 3 | **D-Gluconic Acid** | **Fumaric Acid** | D-Glucosaminic Acid |
| 4 | L-Asn | **Propionic Acid** | Acetoacetic Acid |
| 5 | L-Gln | α-Keto-Glutaric Acid | D-Galactonic Acid-γ-Lactone |
| 6 | L-Ala | Bromo Succinic Acid | Glycolic Acid |
| 7 | L-Pro | **Pyruvic Acid** | D-Galacturonic Acid |
| 8 | Gly-L-Pro | Acetic Acid | **Succinic Acid** |
| 9 | Tyramine | α-Hydroxy Butyric Acid | **Formic Acid** |
| 10 | D-Fructose | L-Lactic Acid | m-Hydroxy Phenyl Acetic Acid |
| 11 | **D-Glucose** | α-Hydroxy Glutaric Acid-γ-Lactone | D-Glucuronic Acid |
| 12 | N-Acetyl-Dglucosamine | α-Keto-Butyric Acid | m-Tartaric Acid |
| 13 | **D-Mannitol** | D-Malic Acid | Mono Methyl Succinate |
| 14 | Glycerol | Methyl Pyruvate | D-Ser |
| 15 | **Itaconic Acid** | L-Ser | **2-Phenylethylamine** |
| 16 | **5-Amino Valeric Acid** | L-Thr | Ammonia |
| 17 | Capric Acid | L-Asp | α-Methyl-D-Glucoside |
| 18 | Sebacic Acid | L-Glu | D-Fructose-6-Phosphate |
| 19 | γ-Amino Butyric Acid | D-Thr | D-Psicose |
| 20 | 4-Hydroxy Benzoic Acid | D-Ala | D-Glucose-6-Phosphate |
| 21 | **ß-Hydroxy Butyric Acid** | D-Asp | **D-Maltose** |
| 22 | Quinic Acid | Gly-L-Glu | L-Fucose |
| 23 | γ -Hydroxy Butyric Acid | 2-Aminoethanol | D-Glucose-1-Phosphate |
| 24 | Caproic Acid | **D-Trehalose** | **D-Xylose** |
| 25 | N-Acetyl-L-Glutamic Acid | Maltotriose | D-Cellobiose |
| 26 | L-PyroglutamicAcid | D-Melibiose | L-Arabinose |
| 27 | **L-Orn** | α-Methyl-D-galactoside | N-Acetyl-ß-D-Mannosamine |
| 28 | Hydroxy-Lproline | Sucrose | Adonitol |
| 29 | L-Ile | Lactulose | D, L-α-Glycerol- Phosphate |
| 30 | L-Arg | α-D-Lactose | Dulcitol |
| 31 | D, L-Octopamine | **D-Ribose** | myo-Inositol |
| 32 | Putrescine | 2-Deoxy Adenosine | D-Sorbitol |
| 33 | **Glucosamine** | Tween 20 | Thymidine |
| 34 | Uric Acid | Tween 40 | α- Keto-Valeric Acid |
| 35 | L-His | Tween 80 | 2- Hydroxy Benzoic Acid |
| 36 | L-Trp | **Butyric Acid** | Oxalic Acid |
| 37 | L-Val | Citramalic Acid | ß-Methyl-D-Glucuronic Acid |
| 38 | Ala-His | Sorbic Acid | Melibionic Acid |
| 39 | Ala-Gln | Succinamic Acid | L-Tartaric Acid |
| 40 | Ala-Glu | Malonic Acid | Citraconic Acid |
| 41 | Gly-Asn | D, L-Carnitine | N-Acetyl-Neuraminic Acid |
| 42 | Gly-Gln | Laminarin | **D-Tartaric Acid** |
| 43 | Ala-Asp | Turanose | L-homoserine |
| 44 | **Ethanolamine** | Xylitol | L- Leu |
| 45 | Agmatine | D-Arabitol | L-Met |
| 46 | Histamine | γ- Amino-N-Butyric Acid | L-Alaninamide |
| 47 | Allantoin | D, L-α-Amino-N-Butyric Acid | L-Phe-Ala |
| 48 | Urea | α-Amino-N-Valeric Acid | Gly-Met |
| 49 | **Acetamide** | ε-Amino-N-Caproic Acid | 3-0-ß-D-Galactopyranosyl-D-Arabinose |
| 50 | **Adenosine** | Parabanic Acid | **Salicylic acid** |
| 51 | **Inosine** | **δ-Amino-N-Valeric Acid** | D-Raffinose |
| 52 | Adenine | N-Acetyl-L-Glutamic Acid | ß-Methyl-D-Xyloside |
| 53 | Guanosine | N-Phthaloyl-L-Glutamic Acid | α-Methyl-D-Mannoside |
| 54 | **Cytosine** | D-Glu | Isoatinose |
| 55 | Cytidine | L-Glu | D-Tagatose |
| 56 | Thymine | L-Lys | Gentiobiose |
| 57 | **Uridine** | L-Tyr | D-Fucose |
| 58 |  | D-Lys | 3-Methyl Glucose |
| 59 |  | D-Asn | D-Ribono-1,4-Lactone |
| 60 |  | L-Gly | L-Glucose |
| 61 |  | Ala-Gly | D-Melezitose |
| 62 |  | Ala-Leu | ß-D-Allose |
| 63 |  | Uracil | D-Arabinose |
| 64 |  | Glucuronamide | ß-Methyl-D-Galactoside |
| 65 |  | N-Acetyl-D-Glucosamine | Amygdalin |
| 66 |  | Nitrite | Sedoheptulosan |
| 67 |  | Nitrate | N-Acetyl-D-Galactosamine |
| 68 |  | Xanthosine | N-Acetyl-D-Glucosaminitol |
| 69 |  | Xanthine | i-Erythritol |
| 70 |  |  | Lactitol |
| 71 |  |  | Maltitol |
| 72 |  |  | D-Val |
| 73 |  |  | **Guanine** |
| 74 |  |  | L- Cys |
| 75 |  |  | L-Citrulline |
| 76 |  |  | Met-Ala |
