## Supplemental Table 2 for "Systematic mapping of chemoreceptor specificities for *Pseudomonas aeruginosa*"

**Table S2 Summary of FRET responses of 16 hybrid chemoreceptors in the presence of 100 μM ligands.** “A”: attractant response; “R”: repellent response; “*”: strong response; “*R or A”*: weak response (or unspecific response); “--”: no response; “blank”: no analysis. The yellow highlights were considered as positive ligands.

| **Categories** | **Compounds** | **Tar** | **PctA** | **PctB** | **PctC** | **PctD** | **CtpM** | **PA2867** | **PctP** | **TlpQ** | **PA1646** | **PA4844** | **PA2573** | **PA1251** | **PA2788** | **PA2920** | **PA5072** | **PA4915** |
| --- | --- | --- | --- | --- | --- | --- | --- | --- | --- | --- | --- | --- | --- | --- | --- | --- | --- | --- |
| **Positive Control** | **D-glucose** | **A*** | **A** | **A** | **A** | **A** | **A** | **A** | **A** | **A** | **A** | **A** | **A** | **A** | **A** | **A** | **A** | **A** |
|  | **Confirmed ligand** | **A*** | **A** | **A** | **A** | **A** | **A** |  |  | **A** |  |  |  |  |  |  |  |  |
| **Organic acids** | **Pyruvate** | **A** | **--** | **--** | ***A*** | **A*** | **A** | ***A*** | ***A*** | **A** | **--** | **A** | **--** | **--** | **--** | **--** | **--** | **--** |
|  | **L-malate** | **A** | **--** | **--** | ***A*** | **--** | **A*** | ***A*** | **A** | **A** | **--** | **A** | **--** | **--** | **A** | **--** | **A** | **A** |
|  | **Itaconate** | **--** | **--** | **--** | **--** | **--** | **--** | **--** | **--** | **--** | **--** | **--** | **--** | **--** | **--** | **--** | **--** | **--** |
|  | **Formate** | **--** | **--** | **--** | **--** | **--** | **--** | **--** | ***A*** | **--** | **--** | **--** | **--** | **--** | **--** | **--** | **--** | **--** |
|  | **Fumarate** | **A** | **--** | **--** | ***A*** | **--** | ***A*** | **A** | ***A*** | **A** | **A** | **A** | **--** | **--** | **--** | **A** | **A** | **A** |
|  | **Propionate** | **--** | **--** | **--** | ***--*** | **--** | **--** | **--** | **--** | **--** | **--** | **--** | **--** | **--** | **--** | **--** | **--** | **--** |
|  | **Succinate** | **A** | **--** | **--** | ***A*** | **--** | **A** | **--** | ***A*** | **A** | **A** | **A** | **--** | **--** | **A** | **--** | **--** | **A** |
|  | **Butyrate** | **--** | **--** | **--** | **--** | **--** | **--** | **--** | **--** | ***A*** | **--** | **--** | **--** | **--** | **--** | **--** | **--** | **--** |
|  | **Glutarate** | **--** | **--** | ***R*** | ***A*** | **--** | **--** | **--** | **--** | **--** | **--** | **--** | **--** | **--** | **--** | **--** | **--** | **--** |
|  | **D-gluconate** | **--** | **--** | **--** | **--** | **--** | **--** | **--** | **--** | **--** | **--** | **--** | **--** | **--** | **--** | **--** | **--** | **--** |
|  | **D-tartrate** | **A** | **--** | **--** | **--** | **--** | **--** | **--** | ***A*** | **--** | **--** | **--** | **--** | **--** | **--** | **--** | **--** | **--** |
|  | **Salicylate** | **--** | **--** | **--** | **--** | **--** | **--** | **R*** | **--** | **--** | **--** | **--** | **--** | **--** | **--** | **--** | **A** | **--** |
|  | **Nicotinate** | **--** | ***A*** | **--** | **--** | ***A*** | **--** | **--** | **--** | **--** | **--** | **--** | **--** | **--** | **--** | **--** | **--** | **--** |
|  | **Isovalerate** | **--** | **--** | **--** | **--** | **--** | **--** | **--** | ***A*** | **--** | **--** | **--** | **--** | **--** | **--** | **--** | **--** | **--** |
|  | **5-Aminovalerate** | **--** | **--** | **--** | **A*** | **--** | **--** | **--** | **--** | **--** | **--** | **--** | **--** | **--** | **--** | **--** | **--** | **--** |
|  | **Methyl 4-aminobutyrate** | **--** | **--** | **--** | **A*** | **--** | **--** | **--** | **--** | **--** | **--** | **--** | **--** | **--** | **--** | **--** | **--** | **--** |
|  | **β-hydroxybutyrate** | **--** | **--** | **--** | **--** | **--** | **--** | **--** | **--** | **--** | **--** | **--** | **--** | **--** | **--** | **--** | **--** | **--** |
|  | **Palmitate** | **--** | ***R*** | **--** | **--** | ***R*^a^** | **--** | ***R*^a^** | **--** | **--** | **--** | **--** | ***R*^a^** | **--** | ***A*^a^** | **--** | **--** | **--** |
|  | **Indoleacetate** | **--** | **--** | **--** | **--** | **--** | **--** | **--** | **--** | **--** | **--** | **--** | **--** | **--** | **--** | **--** | **--** | **--** |
| **Sugars** | **D-fructose** | **A** | **--** | **A** | **--** | **--** | **--** | **--** | ***A*** | **--** | **--** | **--** | **--** | **--** | **--** | **--** | **--** | **--** |
|  | **D-mannitol** | **A*** | ***A*** | ***A*** | **A** | **A** | **A** | **A** | **A** | **A** | **A** | **A** | **A** | **--** | **A** | **A** | **A** | **A** |
|  | **D-xylose** | **--** | **--** | **--** | **--** | **--** | **--** | **--** | **--** | **--** | **--** | **--** | **--** | **--** | **--** | **--** | **--** | **--** |
|  | **D-maltose** | **A*** | **--** | **--** | **--** | **--** | **--** | **--** | **--** | **--** | **--** | **--** | **--** | **--** | **--** | **--** | **--** | **--** |
|  | **D-trehalose** | ***A*** | **--** | **--** | ***--*** | **--** | ***A*** | **--** | ***A*** | ***A*** | **A** | **A** | **A** | **--** | **A** | **A*** | **--** | **--** |
|  | **Glucosamine** | **A*** | **A** | **A*** | **A*** | ***A*** | **A** | **A** | **A*** | ***A*** | ***A*** | **A** | **A** | **A** | **A** | **A** | **A** | **A** |
| **Nucleic acid derivatives** | **Uridine** | ***A*** | ***A*** | ***A*** | **--** | **A** | **--** | **A** | ***A*** | ***A*** | ***R*** | ***A*** | **--** | **--** | **A** | **--** | **--** | **--** |
|  | **D-ribose** | **--** | **--** | **--** | **--** | **--** | **--** | **--** | **--** | **--** | **--** | **--** | **--** | **--** | **--** | **--** | **--** | **--** |
|  | **Cytosine** | **--** | **--** | **--** | **--** | **--** | **--** | **--** | **--** | **--** | **--** | **--** | **--** | **--** | **--** | **--** | **--** | **--** |
|  | **Guanine** | ***R*** | **--** | **A** | ***R*^a^** | **--** | **--** | **--** | **A*** | **--** | **--** | **--** | **--** | **--** | **--** | **--** | **--** | **--** |
|  | **Inosine** | **--** | **--** | **--** | **--** | **--** | **--** | **--** | **A*** | **--** | **--** | **--** | **--** | **--** | **--** | **--** | **--** | **--** |
|  | **Adenine** | **--** | **--** | **--** | **--** | **--** | **--** | **--** | **A*** | **--** | **--** | **--** | **--** | **--** | **--** | **--** | **--** | **--** |
| **Others** | **L-ornithine** | **--** | **A*** | **A*** | **--** | **--** | **--** | **--** | **--** | **--** | **--** | **--** | **--** | **--** | **--** | **--** | **--** | **--** |
|  | **Ethanolamine** | **--** | **--** | **--** | **--** | **--** | **--** | **--** | **--** | **--** | **--** | **--** | **--** | **--** | **--** | **--** | **--** | **--** |
|  | **2-Phenylethylamine** | **--** | **--** | **--** | **--** | **--** | **--** | **--** | ***R*** | **--** | **--** | **--** | **--** | **--** | **--** | **--** | **--** | **--** |
|  | **Acetamide** | **--** | **--** | **--** | **--** | **--** | **--** | **--** | **--** | **--** | **--** | **--** | **--** | **--** | **--** | **--** | **--** | **--** |
|  | **Caffeine** | **--** | **--** | **--** | **--** | **--** | **--** | **--** | **--** | **--** | **--** | **--** | **--** | **--** | **--** | **--** | **--** | **--** |
| **pH** | **pH6.0** | **A*** | **R*** | **R*** | **A*** | **--** | **--** | **--** | **R*** | **R*** | **A*** | **--** | **--** | **--** | **--** | **A** | **--** | **--** |
|  | **pH8.0** | **R*** | **R*** | **A*** | **R*** | **--** | **--** | **--** | **A*** | **R** | **--** | **--** | **R** | **--** | **--** | **--** | **--** | **--** |

**^a^**These ligands were taken as unspecific ligands since the responses were stimulated by their solvents.
