## Supplemental Table 3 for "Systematic mapping of chemoreceptor specificities for *Pseudomonas aeruginosa*"

**Table S3 Strains, plasmids and oligonucleotides used in this study.**

| **Strains and plasmids** | **Genotype or relevant characteristics^a^** | **Reference** |
| --- | --- | --- |
| **Strains** |  |  |
| *Escherichia coli* BL21(DE3) | F^–^ *ompT* *gal* *dcm* *lon* *hsdS_B_*(*r_B_*^–^*m_B_*^–^) λ (DE3 [*lacI* *lacUV5*-*T7p07* *ind1* *sam7* *nin5*]) [*malB*^+^] _K-12_(λ^S^) | (1) |
| *E. coli* DH5α | F^–^ *endA1* *glnV44* *thi-1* *recA1* *relA1* *gyrA96 deoR* *nupG* *purB20* φ80d*lacZ*ΔM15 Δ(*lacZYA-argF*) U169, hsdR17(*r_K_*^–^*m_K_*^+^), λ^–^ | (2) |
| *E. coli* UU1250 | Derivative of RP437; Δ*aer*Δ*tsr*Δ(*tar-tap*) Δ*trg* | (3) |
| *E. coli* VS181 | Derivative of RP437; Δ(*cheYcheZ*)Δ*aer*Δ*tsr*Δ(*tar-tap*) Δ*trg* | (4) |
| *E. coli* CC118λpir | *araD* Δ(*ara*, *leu*) Δ*lacZ74 phoA20 galK thi-1 rspE rpoB argE recA1* λ*pir* | (5) |
| *Pseudomonas aeruginosa* PAO1  (WT-Hiroshima) | Prototroph, FP− (sex factor minus) | (6) |
| *tlpQ::Km* | PAO1 derivative, *PA2654*::Km; Km^R^ | (8) |
| *pctA::Km* | PAO1 derivative, *PA4309*::Km; Km^R^ | (9) |
| *pctB::Km* | PAO1 derivative, *PA4310*::Km; Km^R^ | (10) |
| *pctC::Km* | PAO1 derivative, *PA4307*::Km; Km^R^ | (10) |
| *Pseudomonas aeruginosa* PAO1  (WT-Washington) | Wild type strain | (7) |
| *PA1251::ISlacZ* | PAO1 derivative, *PA1251::ISlacZ*; Tc^R^ | (7) |
| *PA1646**::ISphoA* | PAO1 derivative, *PA1646::ISphoA*; Tc^R^ | (7) |
| *PA2867**::ISlacZ* | PAO1 derivative, *PA2867::ISlacZ*; Tc^R^ | (7) |
| *PA2920**::ISlacZ* | PAO1 derivative, *PA2961::ISlacZ*; Tc^R^ | (7) |
| *PA4290**::ISphoA* | PAO1 derivative, *PA4290::ISphoA*; Tc^R^ | (7) |
| *PA4520**::ISphoA* | PAO1 derivative, *PA4520::ISphoA*; Tc^R^ | (7) |
| *PA4915**::ISphoA* | PAO1 derivative, *PA4915::ISphoA*; Tc^R^ | (7) |
| *mcpA::ISphoA* | PAO1 derivative, *PA0180::ISphoA*; Tc^R^ | (7) |
| *mcpS::ISlacZ* | PAO1 derivative, *PA1930::ISlacZ*; Tc^R^ | (7) |
| *ctpH::ISlacZ* | PAO1 derivative, *PA2561::ISlacZ*; Tc^R^ | (7) |
| *ctpM::ISlacZ* | PAO1 derivative, *PA2652::ISlacZ*; Tc^R^ | (7) |
| *mcpN::ISlacZ* | PAO1 derivative, *PA2788::ISlacZ*; Tc^R^ | (7) |
| *pctD::ISlacZ* | PAO1 derivative, *PA4633::ISlacZ*; Tc^R^ | (7) |
| *ctpL::ISphoA* | PAO1 derivative, *PA4844::ISphoA*; Tc^R^ | (7) |
| Δ*McpK* | PAO1 derivative, Δ*PA5072* | (11) |
| *pctP::Sm* | PAO1 derivative, *PA1608*::Sm; Sm^R^ | This study |
| **Plasmids** | | |
| pET28a (+) | Km^R^; Protein expression vector | Novagen |
| pKG116 | Cm^R^; Expression vector, salicylate inducible; for generation Type 2 and Type 3 hybrid chemoreceptor | (12) |
| pKNG101 | Sm^R^; OriR6K mob, mutant generation vector | (13) |
| pGEMT® | Ap^R^; mutant generation vector | Invitrogen |
| pHC8 | Ap^R^; Expression vector, salicylate inducible; for generation type 1 hybrid chemoreceptor | This study |
| pSB13 | Cm^R^; Tar expression plasmid, pKG116 derivative, T768 was mutated to C768 to remove the NdeI restriction site | (14) |
| pVS88 | Ap^R^; CheY-EYFP / CheZ-ECFP expression plasmid used for type 2 and type 3 hybrid chemoreceptor | (4) |
| pWX16 | Cm^R^; CheY-EYFP / CheZ-ECFP expression plasmid  used for type 1 hybrid chemoreceptor | This study |
| pOB30 | Ap^R^; GFP expression plasmid | (15) |
| pET28_PA2654-LBD | Km^R^; pET28a (+) derivative containing a DNA fragment encoding PA2654-LBD | This study |
| pET28_PA4915-LBD | Km^R^; pET28a (+) derivative containing a DNA fragment encoding PA4915-LBD | This study |
| pET28_PA4844-LBD | Km^R^; pET28a (+) derivative containing a DNA fragment encoding PA4844-LBD | This study |
| pET28_PA5072-LBD | Km^R^; pET28a (+) derivative containing a DNA fragment encoding PA5072-LBD | This study |
| pET28_PA2788-LBD | Km^R^; pET28a (+) derivative containing a DNA fragment encoding PA2788-LBD | This study |
| pET28_PA4310-LBD | Km^R^; pET28a (+) derivative containing a DNA fragment encoding PA4310-LBD | This study |
| pET28_PA4309-LBD | Km^R^; pET28a (+) derivative containing a DNA fragment encoding PA4309-LBD | This study |
| pET28_PA4307-LBD | Km^R^; pET28a (+) derivative containing a DNA fragment encoding PA4307-LBD | This study |
| pET28_PA4633-LBD | Km^R^; pET28a (+) derivative containing a DNA fragment encoding PA4633-LBD | This study |
| pET28_PA1608-LBD | Km^R^; pET28a (+) derivative containing a DNA fragment encoding PA1608-LBD | This study |
| pET28_PA2573-LBD | Km^R^; pET28a (+) derivative containing a DNA fragment encoding PA2573-LBD | This study |
| pET28_PA2652-LBD | Km^R^; pET28a (+) derivative containing a DNA fragment encoding PA2652-LBD | This study |
| pET28_PA2867-LBD | Km^R^; pET28a (+) derivative containing a DNA fragment encoding PA2867-LBD | This study |
| pET28_PA2920-LBD | Km^R^; pET28a (+) derivative containing a DNA fragment encoding PA2920-LBD | This study |
| pET28_PA1251-LBD | Km^R^; pET28a (+) derivative containing a DNA fragment encoding PA1251-LBD | This study |
| pET28_PA1646-LBD | Km^R^; pET28a (+) derivative containing a DNA fragment encoding PA1646-LBD | This study |
| pET28_PA2561-LBD | Km^R^; pET28a (+) derivative containing a DNA fragment encoding PA2561-LBD | This study |
| pET28_PA4520-LBD | Km^R^; pET28a (+) derivative containing a DNA fragment encoding PA4520-LBD | This study |
| PA2654-Tar_T1 | Ap^R^; pHC8 derivative containing a DNA fragment encoding PA2654 [1-373]-Tar [198-553]; | This study |
| PA4915-Tar_T1 | Ap^R^; pHC8 derivative containing a DNA fragment encoding PA4915 [1-198]-Tar [198-553]; | This study |
| PA4844-Tar_T1 | Ap^R^; pHC8 derivative containing a DNA fragment encoding PA4844 [1-312]-Tar [198-553]; | This study |
| PA2573-Tar_T1 | Ap^R^; pHC8 derivative containing a DNA fragment encoding PA2573 [1-190]-Tar [198-553]; | This study |
| PA2788-Tar_T1 | Ap^R^; pHC8 derivative containing a DNA fragment encoding PA2788 [1-187]-Tar [198-553]; | This study |
| PA4310-Tar_T1 | Ap^R^; pHC8 derivative containing a DNA fragment encoding PA4310 [1-288]-Tar [198-553]; | This study |
| PA4310-Tar_T2 | Cm^R^; pKG116 derivative containing a DNA fragment encoding PA4310 [1-353]-Tar [268-553]; | (16) |
| PA4309-Tar_T1 | Ap^R^; pHC8 derivative containing a DNA fragment encoding PA4309 [1-288]-Tar [198-553]; | This study |
| PA4309-Tar_T2 | Cm^R^; pKG116 derivative containing a DNA fragment encoding PA4309 [1-354]-Tar [269-553]; | (16) |
| PA4307-Tar_T1 | Ap^R^; pHC8 derivative containing a DNA fragment encoding PA4307 [1-291]-Tar [198-553]; | This study |
| PA4307-Tar_T2 | Cm^R^; pKG116 derivative containing a DNA fragment encoding PA4307 [1-356]-Tar [268-553]; | (17) |
| PA4633-Tar_T1 | Ap^R^; pHC8 derivative containing a DNA fragment encoding PA4633 [1-369]-Tar [198-553]; | This study |
| PA4633-Tar_T2 | Cm^R^; pKG116 derivative containing a DNA fragment encoding PA4633 [1-435]-Tar [267-553]; | (18) |
| PA1608-Tar_T1 | Ap^R^; pHC8 derivative containing a DNA fragment encoding PA1608 [1-187]-Tar [198-553]; | This study |
| PA1608-Tar_T2 | Cm^R^; pKG116 derivative containing a DNA fragment encoding PA1608 [1-264]-Tar [267-553]; | This study |
| PA2652-Tar_T1 | Ap^R^; pHC8 derivative containing a DNA fragment encoding PA2652 [1-220]-Tar [198-553]; | This study |
| PA2652-Tar_T2 | Cm^R^; pKG116 derivative containing a DNA fragment encoding PA2652 [1-284]-Tar [267-553]; | This study |
| PA5072-Tar_T1 | Ap^R^; pHC8 derivative containing a DNA fragment encoding PA5072 [1-299]-Tar [198-553]; | This study |
| PA5072-Tar_T3 | Cm^R^; pKG116 derivative containing a DNA fragment encoding PA5072 [1-299]-VLYHC-Tar [203-553]; | This study |
| PA2867-Tar_T1 | Ap^R^; pHC8 derivative containing a DNA fragment encoding PA2867 [1-149]-Tar [198-553]; | This study |
| PA2867-Tar_T3 | Cm^R^; pKG116 derivative containing a DNA fragment encoding PA2867 [1-149]-PTTTA-Tar [203-553]; | This study |
| PA2920-Tar_T1 | Ap^R^; pHC8 derivative containing a DNA fragment encoding PA2920 [1-203]-Tar [198-553]; | This study |
| PA2920-Tar_T3 | Cm^R^; pKG116 derivative containing a DNA fragment encoding PA2920 [1-203]-CLAPP-Tar [203-553]; | This study |
| PA1251-Tar_T1 | Ap^R^; pHC8 derivative containing a DNA fragment encoding PA1251 [1-197]-Tar [198-553]; | This study |
| PA1251-Tar_T3 | Cm^R^; pKG116 derivative containing a DNA fragment encoding PA1251 [1-197]-IWSLA-Tar [203-553]; | This study |
| PA1646-Tar_T1 | Ap^R^; pHC8 derivative containing a DNA fragment encoding PA1646 [1-313]-Tar [198-553]; | This study |
| PA1646-Tar_T3 | Cm^R^; pKG116 derivative containing a DNA fragment encoding PA1646 [1-313]-CLYMA-Tar [203-553]; | This study |
| PA2561-Tar_T1 | Ap^R^; pHC8 derivative containing a DNA fragment encoding PA2561 [1-224]-Tar [198-553]; | This study |
| PA2561-Tar_T3 | Cm^R^; pKG116 derivative containing a DNA fragment encoding PA2561 [1-224]-NYKNG-Tar [203-553]; | This study |
| PA4520-Tar_T1 | Ap^R^; pHC8 derivative containing a DNA fragment encoding PA4520 [1-331]-Tar [198-553]; | This study |
| PA4520-Tar_T3 | Cm^R^; pKG116 derivative containing a DNA fragment encoding PA2520 [1-331]-LICDS-Tar [203-553]; | This study |
| pGEMT®-PA1608 | Ap^R^; pGEMT® derivative containing DNA fragment of PA1608 | This study |
| pKNG101-PA1608 | Sm^R^, pKNG101 derivative containing DNA fragment of PA1608 | This study |

^a^Ap, ampicillin; Km, kanamycin; Tc, tetracycline; Cm, chloramphenicol; Sm, streptomycin

| **Oligonucleotide** | **Sequence (5’-3’)** | **Purpose** |
| --- | --- | --- |
| PA2654-LBD_F | GTGCCGCGCGGCAGCCATATGCAACATTCCAGTGTTCTGGTTAAG | Construction of pET28_PA2654-LBD |
| PA2654-LBD_R | ACGGAGCTCGAATTCGGATCCCTATTCGCGGCGCATGTCATCAAGT |  |
| PA4915-LBD_F | GTGCCGCGCGGCAGCCATATGTCTCGTCTGGACGCAAGCTT | Construction of pET28_PA4915-LBD |
| PA4915-LBD_R | ACGGAGCTCGAATTCGGATCCCTAGCGCATACTATCGTAGCGTTT |  |
| PA4844-LBD_F | GTGCCGCGCGGCAGCCATATGCGCGCCATGGAACGTCCGT | Construction of pET28_PA4844-LBD |
| PA4844-LBD_R | ACGGAGCTCGAATTCGGATCCCTACTGCCCTTGCAAACGCTGAC |  |
| PA2573-LBD_F | GTGCCGCGCGGCAGCCATATGCGCCTTGGGAAAGCATTAAGTG | Construction of pET28_PA2573-LBD |
| PA2573-LBD_R | ACGGAGCTCGAATTCGGATCCCTAACGCATGCTATCTTCACGCGC |  |
| PA2788-LBD_F | GTGCCGCGCGGCAGCCATATGAGTATGAGCATCTCGCCGGAAA | Construction of pET28_PA2788-LBD |
| PA2788-LBD_R | ACGGAGCTCGAATTCGGATCCCTACTGAGTATGCTGAACAGAATCC |  |
| PA4310-LBD_F | GTGCCGCGCGGCAGCCATATGAATGATAGCTTACAGCGTGCTT | Construction of pET28_PA4310-LBD |
| PA4310-LBD_R | ACGGAGCTCGAATTCGGATCCCTAGCTAGTGCGCAGTTTGGTT |  |
| PA4309-LBD_F | GTGCCGCGCGGCAGCCATATGAACGACTACTTACAGCGCAAC | Construction of pET28_PA4309-LBD |
| PA4309-LBD_R | ACGGAGCTCGAATTCGGATCCCTAGCTAACGCGAAATTTTGACAGC |  |
| PA4307-LBD_F | GTGCCGCGCGGCAGCCATATGGACTACCGTCAGCGTGAAG | Construction of pET28_PA4307-LBD |
| PA4307-LBD_R | ACGGAGCTCGAATTCGGATCCCTACGAGGTACGAAACTCTGAG |  |
| PA4633-LBD_F | GTGCCGCGCGGCAGCCATATGCGCACCCAAGAACTGGTTC | Construction of pET28_PA4633-LBD |
| PA4633-LBD_R | ACGGAGCTCGAATTCGGATCCCTAACTCATCCCCAGAATATCCTG |  |
| PA1608-LBD_F | GTGCCGCGCGGCAGCCATATGCGCATGGGCAGCCTGAACGA | Construction of pET28_PA1608-LBD |
| PA1608-LBD_R | ACGGAGCTCGAATTCGGATCCCTAGCTGTGGTATACTTCATCA |  |
| PA2652-LBD_F | GTGCCGCGCGGCAGCCATATGAAGAAGCAGGCCGATGCCGA | Construction of pET28_PA2652-LBD |
| PA2652-LBD_R | ACGGAGCTCGAATTCGGATCCCTAGGTGCCGATGCGCTCGTCAAT |  |
| PA5072-LBD_F | GTGCCGCGCGGCAGCCATATGAGCAATCGTACCTTAACCCATC | Construction of pET28_PA5072-LBD |
| PA5072-LBD_R | ACGGAGCTCGAATTCGGATCCCTAACGATCACGGGACGCGGTA |  |
| PA2867-LBD_F | GTGCCGCGCGGCAGCCATATGCAACAAGAAGCCGCGGGG | Construction of pET28_PA2867-LBD |
| PA2867-LBD_R | ACGGAGCTCGAATTCGGATCCCTAACGATCTGCAAACACTTGCCAA |  |
| PA2920-LBD_F | GTGCCGCGCGGCAGCCATATGCAATTAAATGAAAGCAATCAACG | Construction of pET28_PA2920-LBD |
| PA2920-LBD_R | ACGGAGCTCGAATTCGGATCCCTACTGCTGCCAACGATGGC |  |
| PA1251-LBD_F | GTGCCGCGCGGCAGCCATATGCGCATGGGTAGCCTGAATCG | Construction of pET28_PA1251-LBD |
| PA1251-LBD_R | ACGGAGCTCGAATTCGGATCCCTAATCTGCCGCCTCAGCTGTAGA |  |
| PA1646-LBD_F | GTGCCGCGCGGCAGCCATATGCAACGCTTTGCCCGCTTACA | Construction of pET28_PA1646-LBD |
| PA1646-LBD_R | ACGGAGCTCGAATTCGGATCCCTACGAATTTGTGCGCAGGGAGTG |  |
| PA2561-LBD_F | GTGCCGCGCGGCAGCCATATGACCGGTGATGTGCGTGCATAT | Construction of pET28_PA2561-LBD |
| PA2561-LBD_R | ACGGAGCTCGAATTCGGATCCCTAGGTGCGACGCGCCTCAGCGCT |  |
| PA4520-LBD_F | GTGCCGCGCGGCAGCCATATGTATCTGGTACGTGACGCATAC | Construction of pET28_PA4520-LBD |
| PA4520-LBD_R | ACGGAGCTCGAATTCGGATCCCTACTGCGTACGGACCTGTTGT |  |
| PA1608_MUT_F | AGTTTCGCCCTGATCACCG | Construction of pKNG101-PA1608 |
| PA1608_MUT_R | TTCCTGGATGTTGGCGATCAT |  |
| pKG116-seq_F | AAGCCATAAGGAGTACCATATG | Sequencing for hybrid  Chemoreceptor_T2&T3 |
| pKG116-seq_R | TTACTTATTTATCCGCGGATC |  |
| pET28a-seq_F | GTGCCGCGCGGCAGCCATATG | Sequencing for LBD expression plasmids |
| pET28a-seq_R | ACGGAGCTCGAATTCGGATCCCTA |  |
| Hybrid-Seq­­­_F | AGCCATAAGGAGTACCATATG | Sequencing for hybrid  Chemoreceptor_T1 |
| Hybrid-Seq­­­_R | AGCAGAATCAATACCACCAC |  |
| PA1608-T2_F1 | AGCCATAAGGAGTACCATATGATGTCTTTGCGCAGTATGCC | Construction of PA1608-Tar_T2 |
| PA1608-T2_R1 | CCTTCGCGGACATGCTGGATGGTCTGGCGCAGTTG |  |
| PA1608-T2_F2 | CATCCAGCATGTCCGCGAAGGTTCAGAT |  |
| Tar-T2_R2 | GGATCCTCAAAATGTTTCCCAGTTT |  |
| PA2652-T2_F1 | AGCCATAAGGAGTACCATATGATGCGTCTGACCCTGAA | Construction of PA2652-Tar_T2 |
| PA2652-T2_R1 | CCTTCGCGGACATGCCGCACCAGACCGTGGATC |  |
| PA2652-T2_F2 | GTGCGGCATGTCCGCGAAGGTTCAGAT |  |
| Tar-T3_F | TTGATTCTGCTGGTGGCGTG | Construction for hybrid  Chemoreceptor_T3 |
| pKG116_Tar-T3_R | TACTTATTTATCCGCGGATCCTCAAAATGTTTCCCAGTTTGGATCTTG |  |
| PA5072-T3_F | AGCCATAAGGAGTACCATATGATGTATGATTGGTGGGTCTTG | Construction of PA5072-Tar_T3 |
| PA5072-T3_R | CCACCAGCAGAATCAANNNNNNNNNNNNNNNGGCTGCAATCAGCCATAA |  |
| PA2867-T3_F | AGCCATAAGGAGTACCATATGATGGGTACGTGGATTTCGGAT | Construction of PA2867-Tar_T3 |
| PA2867-T3_R | CCACCAGCAGAATCAANNNNNNNNNNNNNNNAAGCATCAGCACTAAAAC |  |
| PA2920-T3_F | AGCCATAAGGAGTACCATATGATGCTGCAATGGTTCGCC | Construction of PA2920-Tar_T3 |
| PA2920-T3_R | CCACCAGCAGAATCAANNNNNNNNNNNNNNNCAACAGTCCAAAGGCAAC |  |
| PA1251-V3_F | AGCCATAAGGAGTACCATATGATGCTTCTTCGTCGCATT | Construction of PA1251-Tar_T3 |
| PA1251-T3_R | CCACCAGCAGAATCAANNNNNNNNNNNNNNNTAACAACACTGCCAGAATG |  |
| PA1646-T3_F | AGCCATAAGGAGTACCATATGATGTTGGGATTATTACGTCG | Construction of PA1646-Tar_T3 |
| PA1646-T3_R | CCACCAGCAGAATCAANNNNNNNNNNNNNNNAACCAGCAATGCTAAAAC |  |
| PA2561-T3_F | AGCCATAAGGAGTACCATATGATGCCTGCTTCTCCTGG | Construction of PA2561-Tar_T3 |
| PA2561-T3_R | CCACCAGCAGAATCAANNNNNNNNNNNNNNNACCAATCAGCACCAATGA |  |
| PA4520-T3_F | AGCCATAAGGAGTACCATATGATGAAAACCGTATTGTACCCG | Construction of PA4520-Tar_T3 |
| PA4520-T3_R | CCACCAGCAGAATCAANNNNNNNNNNNNNNNCACCACCGCCAACGCTGC |  |
