## Supplemental Table 4 for "Systematic mapping of chemoreceptor specificities for *Pseudomonas aeruginosa*"

**Table S4 Experimental conditions used for microcalorimetric titrations**

| **Protein** | | **Ligand** | | **Temperature (ºC)** | **Injection volume (µL)** | **Analysis buffer composition** |
| --- | --- | --- | --- | --- | --- | --- |
| **Name** | **Concentration (µM)** | **Name** | **Concentration (mM)** |  |  |  |
| PctA-LBD | 39 | L-ornithine | 1 | 30 | 4.8 | 5 mM Tris, 5 mM Pipes, 5 mM Mes, 10 % glycerol (vol/vol), 150 mM NaCl, pH 7.0 |
| PctB-LBD | 50 | L-ornithine | 20 | 30 | 12.8 | 5 mM Tris, 5 mM Pipes, 5 mM Mes, 10 % glycerol (vol/vol), 150 mM NaCl, pH 7.0 |
| PctC-LBD | 120 | Methyl 4-aminobutyrate | 20 | 25 | 14.4 | 5 mM Tris, 5 mM Pipes, 5 mM Mes, 10 % glycerol (vol/vol), 150 mM NaCl, pH 7.0 |
|  | 120 | 5-aminovalerate | 5 | 25 | 3.2 |  |
| TlpQ-LBD | 93 | 2-phenylethylamine | 20 | 15 | 11.2 | 5 mM Tris, 5 mM Pipes, 5 mM Mes, 10 % glycerol (vol/vol), 150 mM NaCl, pH 7.0 |
|  | 147 | Tyramine | 20 | 25 | 6.4 |  |
| PA1608-LBD | 65 | Hypoxanthine | 2 | 20 | 4.8 | 10 mM sodium phosphate, pH 5.5 |
| PA4915-LBD | 92 | 2-phenylethylamine | 5 | 25 | 12.8 | 40 mM Potassium phosphate, 10 % glycerol (vol/vol), pH 7.0 |
